# Cell surface excess is essential for protrusions and motility in 3D matrix

**DOI:** 10.1101/2022.08.12.503602

**Authors:** Maryna Kapustina, Donna Li, James Zhu, Brittany Wall, Violetta Weinreb, Richard E. Cheney

## Abstract

To facilitate rapid changes in morphology without endangering cell integrity, each cell possesses a substantial amount of cell surface excess (CSE) that can be promptly deployed to cover cell extensions. CSE can be stored in different types of small surface projections such as filopodia, microvilli, and ridges, with rounded bleb-like projections being the most common and rapidly achieved form of storage. We demonstrate in this paper that cells migrating in 3D collagen use CSE to cover the developing protrusions. After retraction of a protrusion, the CSE this produces is stored over the cell body similar to the CSE produced by cell rounding. For the coordinated process of CSE storage and release, all cells should have specific mechanisms of regulation, and we hypothesize that microtubules (MT) play an important role in this mechanism. We show here that different effects of MT depolymerization on cell motility such as inhibiting mesenchymal motility and enhancing amoeboid, can be explained by the essential role of MT in CSE regulation and dynamics.

## Introduction

Crucial cell functions including migration, cytokinesis, and differentiation involve major changes in cell shape. Morphological transformations often result in dramatically different cell shapes, requiring cells to develop an efficient mechanism to manipulate surface area while keeping volume nearly constant. The periphery of the cell, which we term the cell surface, must be very flexible to accommodate fast shape changes during migration while maintaining the integrity of the cell. It is comprised of a plasma membrane (PM) with a thin (~<200-500 nm) layer of cytoskeleton structure known as the cell cortex, which is coupled to the PM by adaptor proteins and consists of an F-actin network cross-linked by actin-binding proteins [1–6]. Importantly, despite the flexibility of the plasma membrane, it can only undergo ~3% areal extension without rupturing [7–10]. Thus, to adapt to diverse environmental requirements, a sufficient reservoir of the plasma membrane that is immediately deployable is critically important for cell survival.

Pioneering work with scanning electron microscopy [11–14] revealed that the surface of rounded cells is covered with numerous small surface projections such as blebs, microvilli, filopodia, and ridges. These surface projections largely disappear upon cell spreading, leading to the hypothesis that they serve as a reservoir for storing the excess of the plasma membrane that can be flattened and reused during surface expansion. In this paper, we define *cell surface excess (CSE)* as the difference between actual cell surface area, which includes the surface area of all small projections, and the area required to cover the cell volume with a smooth surface layer.

Studies from the past decades provided evidence that CSE can be utilized to cover increasing surface areas or to buffer mechanical stresses occurring in diverse physiological processes such as phagocytosis, cellularization in Drosophila embryos, or cell spreading [13–18]. Our study of the spontaneous and microtubule depolymerization induced morphological oscillations in non-spread (rounded) cells with a period on the order of one to a few minutes demonstrated [19, 20] that traveling wave of membrane-cortex density exhibited by these cells originates from the periodic accumulation and release (compression and dilation) of the numerous small surface projections and folds and that an excess of cell surface area is a requirement for this phenomenon. We suggested that this mechanism for rapidly altering cell surface area is essential for many morphological transformations and can provide a model for cell shape transformation during migration. Many cells exhibit rich phenotypic plasticity during locomotion and adapt their mechanisms of migration to fit their 3D tissue microenvironment [21–23]. The two major modes of motility, mesenchymal and amoeboid, have different cytoskeletal, biomechanical, and signaling properties [24–26]. Cells utilizing the mesenchymal mode of motility rely on acto-myosin fibers to produce contractile forces, on strong focal adhesions to the extra-cellular matrix (ECM) to apply traction forces, and on ECM-degrading enzymes to perform ECM remodeling and generate a path for translocation [27]. Amoeboid migration commonly refers to the movement of more rounded cells that lack mature focal adhesions and stress fibers [28–30]. The specific migration style was first observed in amoebae and later found in many other cells, including leukocytes and certain types of tumor cells. The characteristic feature of amoeboid migration is a high level of actomyosin contractility at the cell cortex mostly mediated by Rho signaling. The amoeboid mode of migration allows cells to move at high velocities by adapting their bodies to the pre-existing spaces and squeezing through the gaps in the ECM fibers without developing mature adhesions or inducing ECM degradation [31, 32].

In this paper, we demonstrate that cells in a 3D collagen environment maintain a large amount of CSE stored in small surface projections. We also demonstrate that the process of releasing stored surface by flattening small surface projections is coordinated with the extension of large cell protrusions. Conversely, the process of forming small surface projections, primarily blebs, is coordinated with the retraction of the cell protrusions. We also demonstrate that an intact MT system is necessary for proper CSE regulation and stability. We suggest that despite many differences between mesenchymal and amoeboid motility, both modes of motility rely on the proper regulation of CSE.

## Results

### 1. Rounded cells possess large amounts of cell surface excess

When a spread cell (Figure 1A) is proteolytically detached, it rapidly rounds on a time scale of ~30-60 s with a significant reduction in apparent cell surface area (Figure 1B). We estimated previously [33] that despite limited time for membrane endocytosis, this reduction for CHO cells was 3.5± 2.25 fold and up to 12 fold for Swiss 3T3 cells. Importantly, the fluorescence signal from the cortical F-actin, which is practically unresolved in spread cells [34], becomes visible as a thick, bright band at the margins of rounded cells (Figure 1B,D). The same cells visualized using high-resolution scanning electron microscopy reveal a highly convoluted surface morphology that is capable of storing a large amount of surface area (Figure 1C,E,F). Note that a smooth surface lacking small membrane projections covers only fully spread cells (Figure 1C,G). The morphology of membrane convolutions is characterized by several types of small projections (Figure 1H): short spherical blebs, short and slender microvilli or filopodia-like structures, and ruffles. The typical length of small filopodia-like projections is between 200 nm and 2000 nm, with an average length of 626 nm +−360 nm (n=114) and previously, we estimated that the average radius of a small bleb-like structure on a cell surface is below 250 nm [33]. Transmission electron microscopy images of anti-GFP immunogold staining of rounded CHO cells stably expressing biomarker of filamentous actin, Lifeact-GFP, revealed that all these small projections contain F-actin (Figure 1I,J). These small structures are tightly packed on the cell periphery (see the plot in Figure 1J) and are often unrecognizable as separate structures at optical microscope resolution (Figure 1D), where they appear as a thick homogenous actin layer and are frequently assumed to be a continuous cortical actin network. A simple estimate of the perimeter length on a random position of a rounded cell from a TEM image (Figure 1I) demonstrates that the area with small projections has approximately three-fold longer perimeter than a smooth surface without the projections, highlighting the immense amount of CSE contained in rounded cells.

**Figure 1.**
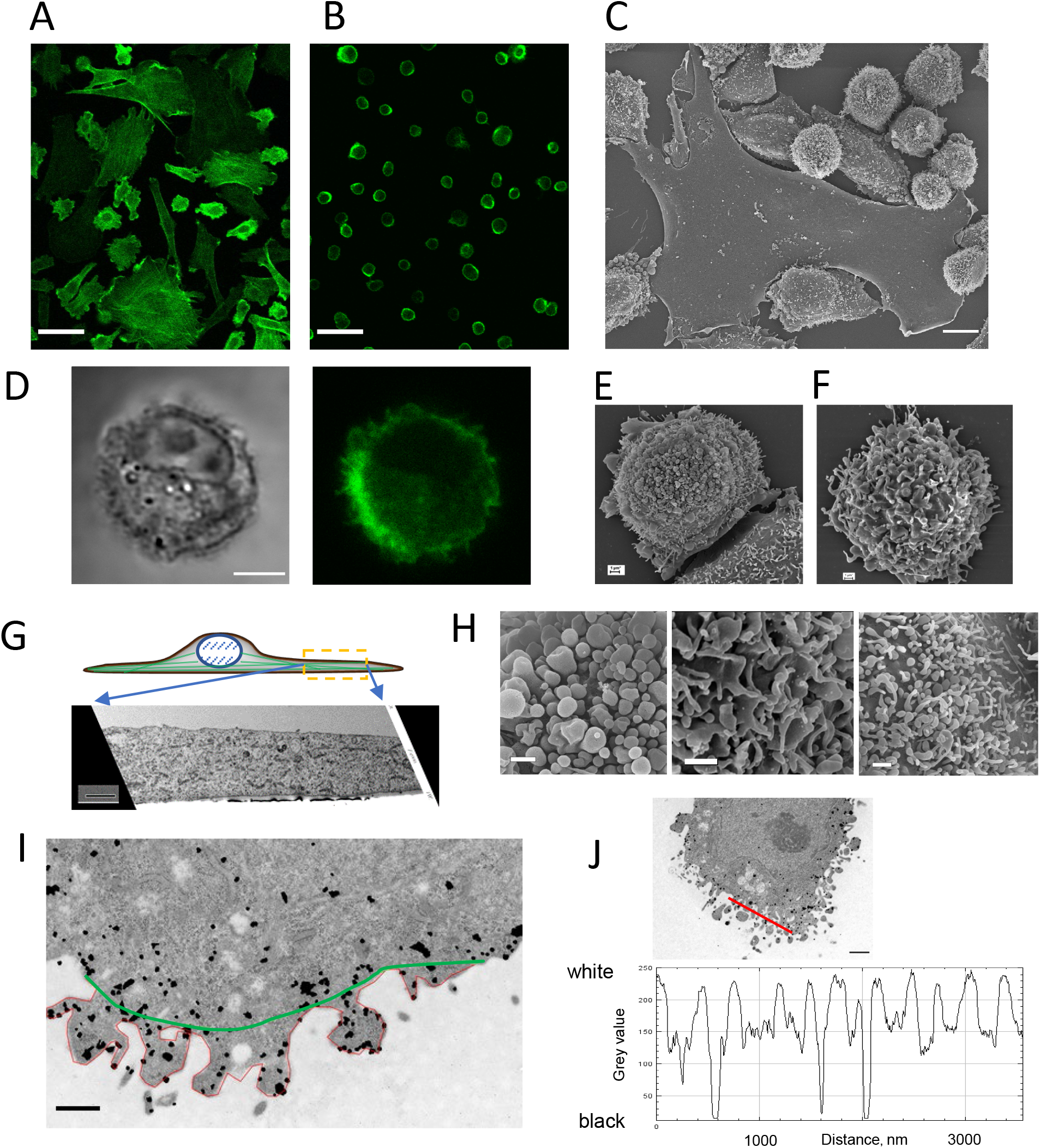
Rounded cells on a 2D substrate possess an ample amount of cell surface excess. **A-B**. The same CHO cells with stable expression of F-actin marker (Lifeact-GFP) were imaged while spread on the glass (A) and immediately after rounding (B). **C**. Scanning electron microscopy (SEM) image demonstrates the difference in the surface morphology between spread and rounded CHO cells. **D.** Images of representative CHO cell in the rounded state (DIC-left and Fluorescence Lifeact-GFP-right) demonstrates the apparent thickness of actin cortex visible at optical microscope resolution. **E,F** Examples of SEM images of CHO (E) and DU-145 (F) cells fixed 20 min after rounding. **G**. Transmission electron microscopy image (TEM) image of fully spread CHO cells sliced perpendicularly to the substrate as shown in the cartoon above the image. The image demonstrates the absence of small membrane projections on the surface. **H.** Magnified view of three different types of small surface structures (blebs, ridges, and microvilli) on the periphery of different rounded cells. **I,J.** TEM image of GFP immunogold staining of rounded CHO cells with stable expression of Lifeact-GFP. Black dots which represent gold particles show the position of actin filaments. I. Green and red lines demonstrate the difference in the length between cell periphery with small protrusions (red) and without (Green) (ratio between the length of two lines is ~3). **J.** Plot of pixels intensity taken along the red line on J. The plot demonstrates that the distance between small surface projections is in the range of 50-200 nm which is below the optical resolution of a regular confocal microscope. Scale bars: A,B = 50 um; C= 10 um; D=5 um; E,F = 1 um; G=2 um; H=1um; I= 500 nm; J=1um

### 2. Electron micrographs demonstrate surface smoothing during protrusion

We hypothesize that a fast-growing cell protrusion can acquire most of its surface from the surface excess stored around the cell body (see the cartoon in Figure 2A). This mechanism would be analogous to the initial stages of cell spreading, where cell surface stored in small surface projections on rounded cells is employed to cover the spreading areas. To find evidence for such a mechanism, we used light microscopy and SEM to visualize the morphology of DU-145 cells (a human metastatic prostate cancer line) during the initial stages of cell attachment to a glass substrate. Light microscopy shows that the rounded DU-145 cells often develop short protrusions (Figure 2B, blue arrows) that connect the cells with the substrate prior to cell spreading. Although these short protrusions can look like elongated blebs, they often persist for over an hour, much longer than the ~1-3 minutes lifetime reported for blebs [35, 36]. SEM images in Figure 2C,F demonstrate that the surface of these short protrusions is relatively smooth (green arrows) and their edges are characterized by elongated narrow filopodia-like or blunted projections that contact the coverslip (red arrows). The blebs which remained at the leading margin of the protrusion (orange arrows) are in a position to provide a surface for further growth. A similar process of blebs transforming into small protrusive structures on the cell periphery is captured in images (Figure 2G,H).

**Figure 2.**
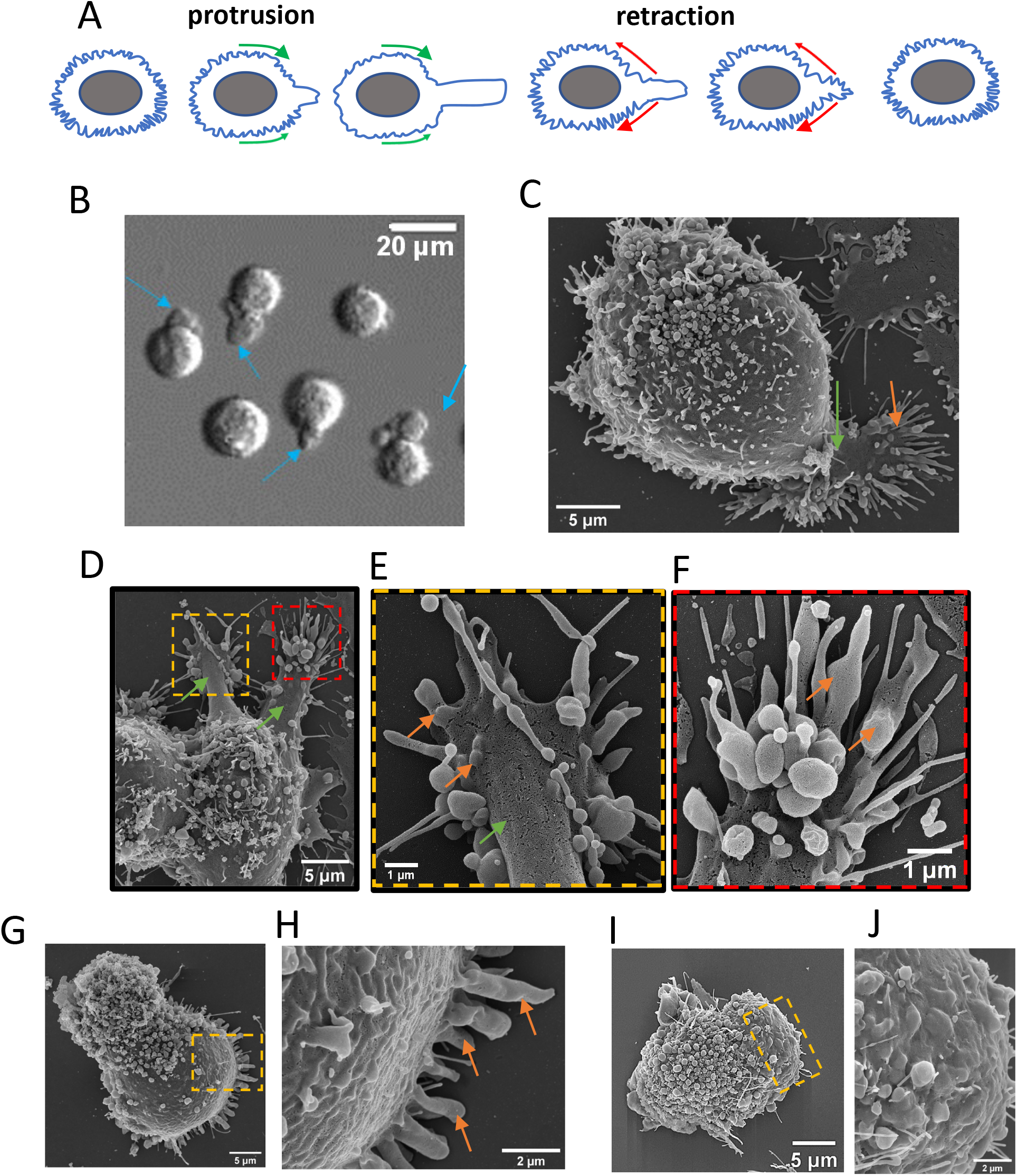
CSE is released during protrusion development and spreading. **A**. Cartoon illustrating the hypothetical process of protrusion development and retraction in correlation with CSE release and accommodation. **B.** DIC image of DU-145 cells 30 min after plating on the glass. Note the short protrusions (blue arrows) that serve as the initial points of attachment. The cell bodies can hang on this “hand” for several hours before spreading occurs. **C-H**. SEM images of DU-145 cells in 30 min after plating on glass. The enlarged parts of the image (D) on E (yellow box) and F (red box) provide several examples of the flat membrane surface on the main protrusion (green arrows) with the blebs in the process of transformation into small protrusive structures (orange arrows). **C, G, H**, Images demonstrate a plasma membrane flattening and possible flow during spreading. One part of the cell is covered with small surface structures while on the opposite part these structures disappear potentially providing the cell surface for spreading. We hypothesize that blebs close to the substrate become elongated (orange arrows) as the cell prepares to extend the periphery. **I-J.** SEM images of representative CHO cell 30 min after plating on glass. Images demonstrate the same process of small surface structure flattening and possible plasma membrane flow during spreading as for the DU-145 cells on (G,H). Scale bar: B = 20 um; C,D,G,I = 5 um; H,J = 2 um; E,F = 1 um.

SEM images in Figure 2C,G also demonstrate that while the cell surface distal to the extending region is covered with small projections holding a large amount of surface excess, these structures become flattened and the cell surface becomes much less convoluted in regions adjacent to the extension. The SEM images of rounded CHO cells revealed similar changes in cell surface morphologies during the initial steps of spreading (Figure 2I,J). Overall, SEM images demonstrate how small surface structures are flattened and the released surface is reused for protrusion development and extension of the cell periphery.

### 3. CHO cells embedded in a 3D matrix preserve surface excess for successful morphological transformations

While cell rounding after detachment from a coverslip clearly demonstrates the phenomenon of CSE, it is not a naturally occurring condition. Therefore, we next explored whether cells maintain CSE under physiologically relevant conditions such as during migration in a 3D collagen matrix.

First, we investigated the morphology and behavior of CHO cells. These cells don’t possess the specific integrin (α2) for binding to collagen I fibers [37, 38]. In a 3D collagen matrix, they maintain either a rounded or a slightly polarized shape with infrequent protrusions (Figure 3A). Visual evaluation confirmed that CHO cells do not modify collagen fiber structure but rather stay and move within the unmodified collagen network (SupFigure 1A). Surprisingly, rounded CHO cells that had been embedded longer than 24h in 3D collagen retained a convoluted surface morphology that can store a large amount of CSE similar to that of newly rounded cells on a 2D substrate. Small surface projections are often clearly visible on the surface of the cells (Figure 3B), and the cortical actin associated with these structures is underlain by a band of myosin II, very similar to CHO cells after rounding on a coverslip (SupFigure 1B). The radius of rounded CHO cells embedded in collagen for more than 24 h was between 6 μm and 9.5 μm with an average of 7.91±1.41 μm (N=128), which is not significantly different from CHO cells after rounding on a 2D substrate (8.07 ±1.62 μm, N=226, p>0.1). These results clearly show that the CSE phenomenon is not an artifact of the experimental conditions but is a part of normal cell physiology and therefore should be regulated by some specific mechanisms.

**Figure 3.**
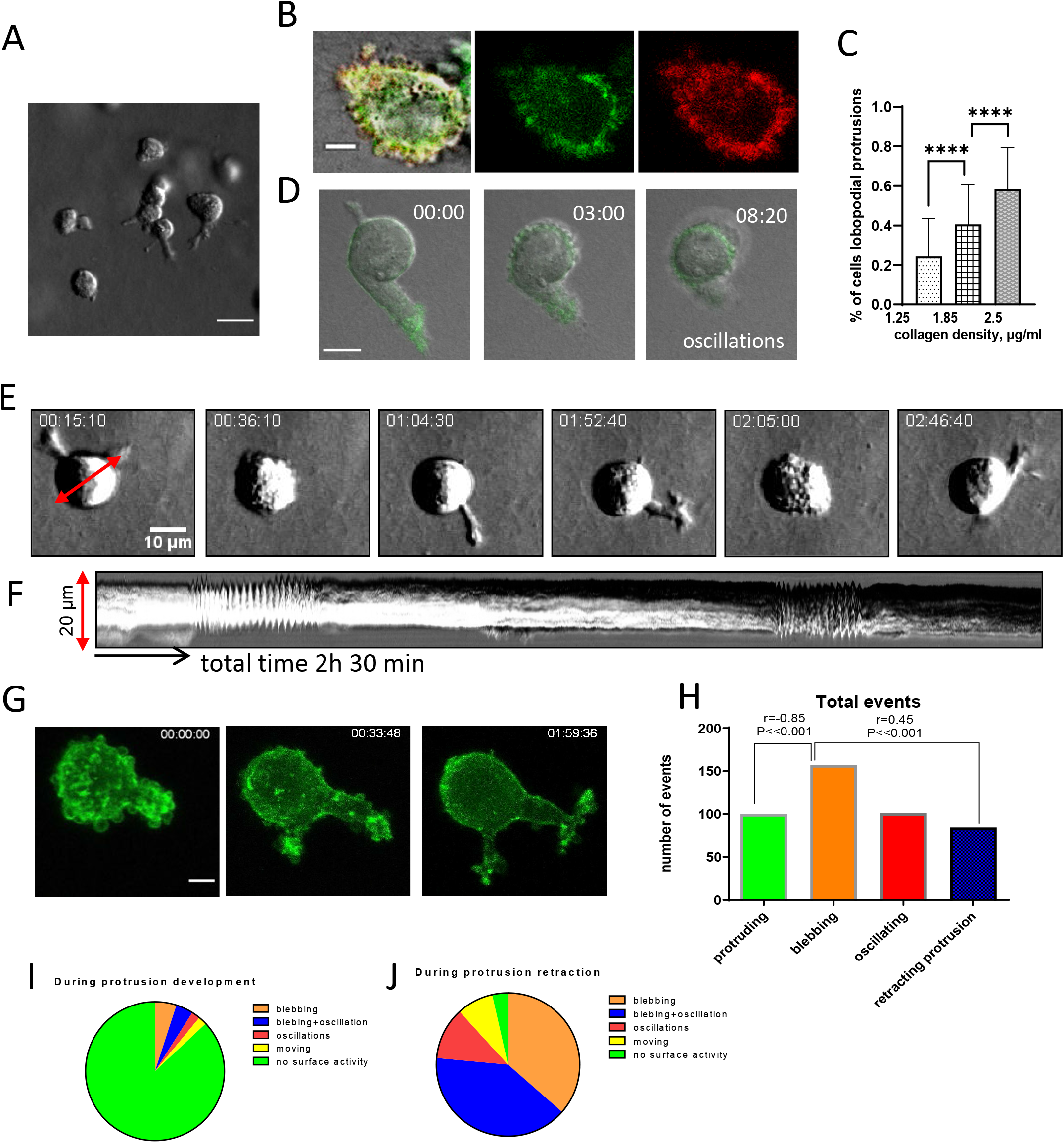
Cells embedded in a 3D matrix preserve surface excess and use it for protrusion development. **A**. DIC image shows different morphology of CHO cells embedded in the 3D collagen for 24h. **B**. Cells embedded in a 3D matrix preserve CSE. Morphology of rounded CHO cell transfected with membrane marker mRFP-PMT and Lifeact-GFP. **C.** The chart presents the proportion (percent) of CHO cells with lobopodial protrusions in collagen with different densities (N=250, 2 experiments). **D.** Merged DIC and F-actin fluorescence images of the CHO cell during protrusion retraction which results in cell blebbing and oscillations (movie 6). Note the smooth cell surface while protrusion is fully extended. **E,F.** DIC image (E) and kymograph F) of CHO cell in collagen during several cycles of protrusion-retraction. (movie 7). **G.** Maximum intensity projection image of CHO (Lifeact-GFP) cell in collagen before and after the development of the protrusion. **H-J.** The charts visualize the results of quantitative analysis of cell protrusive activity in correlation with surface dynamics (n=258, 37cells). Scale bars: A = 20 μm, B,G = 5 μm, C,E = 10 μm.

In the absence of strong collagen adhesions and matrix-degrading proteases, CHO cells can move either by developing blunt cylindrical protrusions <u>(movie 1)</u> similar to lobopodial motility [39] or by using protrusive bulges (blebby type of amoeboid migration, <u>movie 2)</u>. The latter has similarities to the compression-dilation mode seen in oscillatory cells where the combined image of DIC and F-actin (Lifeact-GFP) fluorescence signals shows stretching of the cell membrane-cortex layer during bulge extension (SupFigure 1C-E, <u>movie 3,4</u>). During morphological oscillations, the highly periodic process of stretching (dilation) and compression of the membrane-cortex layer often creates a traveling wave of cortex density around the cell periphery (SupFigure 1D, <u>movie 5)</u> [20].

The same CHO cell can demonstrate both types of <u>protrusions</u> in the collagen: lobopodial and blebby protrusion. Our analysis of CHO cells embedded in collagen matrices with different densities revealed that the cells develop lobopodial protrusions substantially more often in the dense collagen matrix (Figure 3C). This can be explained by the difference in the size of
collagen pores, implying that a slim lobopodial protrusion can penetrate a narrower space between collagen fibers than a large rounded bleb.

To better understand the amount of CSE maintained by CHO cells during migration, we estimated how much surface excess is produced by retraction of the lobopodial protrusion, similar to the one that presented in Figure 3D (SupFigure 2A, <u>movie 6</u>). Assuming that the cell volume is conserved during protrusion retraction, our estimations show that approximately 64% (240 μm^2^) of the protrusion surface in this particular cell is no longer needed for covering cell volume after protrusion retraction and therefore must be stored as CSE or endocytosed. Our analysis of the large population of CHO cells with lobopodial protrusions (103 protrusions on 53 cells, average length of 23.4± 13.5 μm and width 4.94±1.4 μm) shows that between 60% to 80% of protrusion surface rapidly became cell surface excess after protrusion withdrawal.

### 4. CHO cells employ CSE to cover the extension of protrusion while CSE is accumulated during retraction

According to our hypothesis, flattening of pre-existing surface projections to release CSE should be observed in CHO cells during 3D migration. To test this, we imaged CHO cells embedded in collagen matrices for prolonged periods and analyzed their surface and protrusive dynamics. Figure 3E and <u>movie 7</u> demonstrate a protrusive phenotype of CHO cells in collagen matrices where these cells undergo morphological oscillations and blebbing prior to the extension of a lobopodial-like protrusion. Simultaneously with the initiation of a protrusion, the morphological oscillations ceased and the blebbing decreased. After the extension of a large, stable protrusion, the cell exhibited low surface dynamics and only a few blebs. Kymographs (Figure 3F) describing cell surface activity as a function of time, which corresponds to the time steps on the frames of a DIC recording, demonstrate the correspondence in time between cell surface apparent activity and protrusion dynamics. The transition from blebbing to a smooth surface after protrusion growth is clearly visible on confocal fluorescence imaging of F-actin (Figure 3G). In contrast, in the initial stages of protrusion withdrawal, the cell surface became covered with dynamic blebs which were highly pronounced after retraction (Figure 3D and SuplFigure 2D,E, <u>movie 8)</u>. Often, during the later stages of protrusion retraction, morphological oscillations are initiated (Figure 3D,E and SupFigure 2D). The retraction of a protrusion usually leaves a region on the cell periphery densely populated with blebs, which remain local until the excess surface stored in those blebs is redistributed around the cell surface (SupFigure 2D,F) and movie 8 demonstrates how the CSE can be rearranged by the traveling wave of cortex density during the cell oscillations. Blebs and morphological oscillations often characterize the rounded non-polarized cell until a new protrusion emerges.

Using the time-lapse recording of CHO cells (37 cells in 5 experiments) we collected information about changes in cell protrusions, motility, surface blebbing, and morphological oscillations (258 events total). The data analysis (Figure 3H-J) confirmed statistically significant *positive* correlations between cycles of high surface activity (apparent blebbing) and protrusion retraction (r=0.45, P<<0.001). It also showed a highly significant *negative* correlation between protrusion development and surface blebbing (r=-0.85, P<<0.001), confirming our hypothesis that the cell body’s surface relaxes and flattens during lobopodial protrusion development.

### 5. Cells with mesenchymal motility also employ CSE for the protrusion in 3D collagen

Next, we asked if the cells that are known to employ a mesenchymal mode of motility also maintain CSE and use it for protrusion development during 3D motility. To test this, we used several different cell lines. First, we used Walker carcinoma (WC) cells, a rat mammary carcinoma line that is highly motile. The average apparent radius of rounded WC cells on a 2D surface and after embedding into 3D collagen were very similar, R_2D_=6.02±0.65 μm N=53 vs R_3D_=6.04±0.51 μm, N=34. Shortly after embedding in 3D collagen, WC cells developed multiple very long (>100 μm) and thin (0.4 μm −1 μm) protrusions (Figure 4A, <u>movie 9</u>) and were capable of migration speed up to 2 μm/min. The fast migration of these cells involved substantial collagen contraction (Figure 4B, <u>movie 10</u>). Rounded WC cells had a convoluted surface morphology (Figure 4D), covered with blebs and small projections which hold a large amount of CSE, while the surfaces of polarized cells possessing long protrusions were smooth with no apparent blebbing (Figure 4C). Visual and kymographic analysis demonstrate that blebbing on rounded cells decreased simultaneously with protrusion initiation (Figure 4E, <u>movie 11</u>). Blebbing and oscillations return during full or partial protrusion retraction; the blebbing may be global, involving the entire cell surface, or local, only in the area near the retracting protrusion (Figure 4F, <u>movie 12</u>). If a new protrusion started to emerge simultaneously with the withdrawal of an old protrusion, blebbing was often diminished, suggesting that the surface surplus was immediately re-used to allow the extension of the new protrusion (Figure 4G).

**Figure 4.**
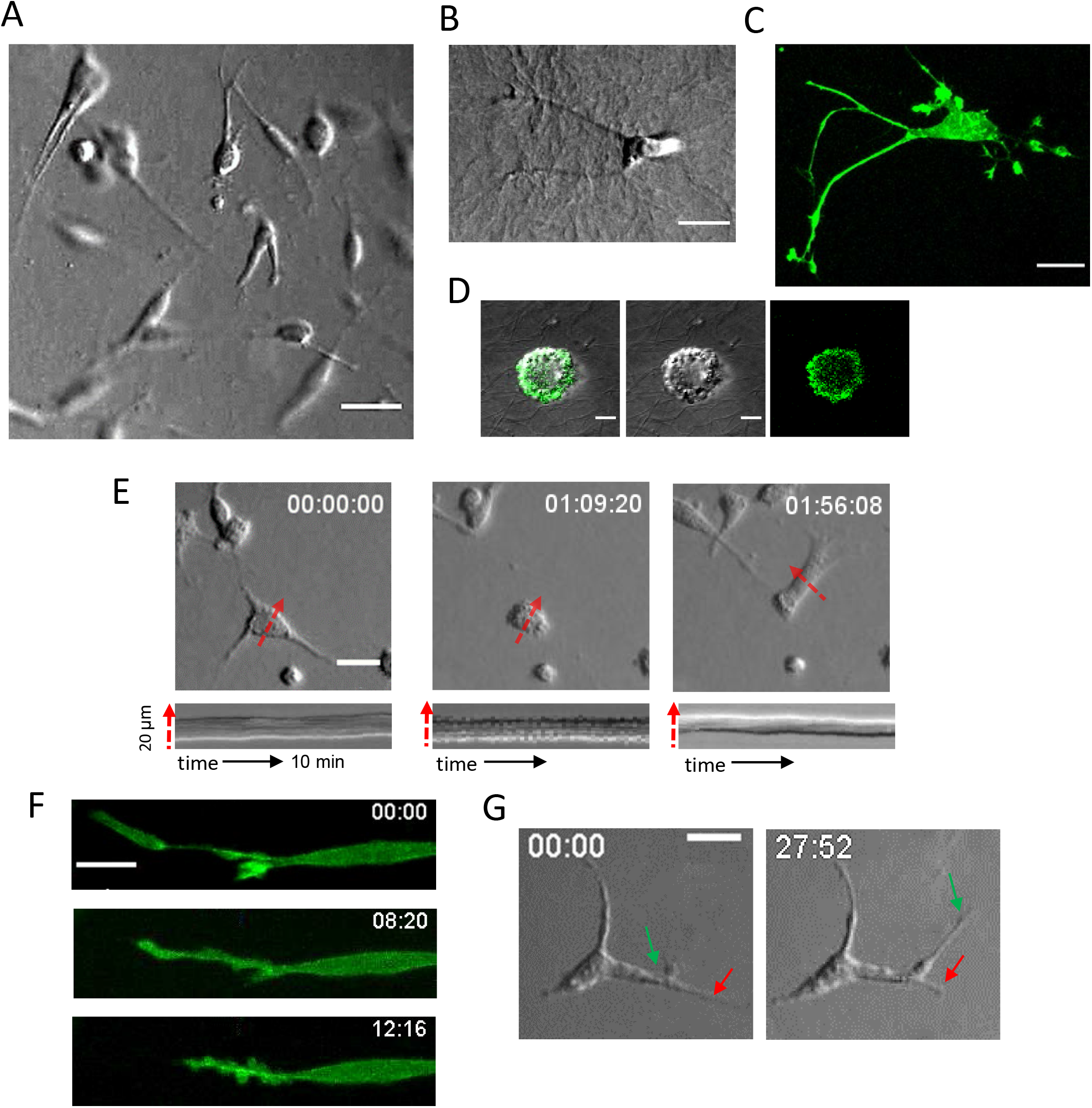
Correlation between protrusion dynamics and apparent blebbing in WC cells. **A-F.** WC cells in 3D collagen. **A.B** DIC recording show WC morphology and strong collagen modification during cell motility (movie 9,10). **C.** Maximum intensity projection image of representative WC cell with Lifeact-GFP signal. The MIP image was built from Z-stack images taking each 1μm through 47 μm of collagen thickness. **D.** Merged, DIC, and fluorescence (Lifeact-GFP) images of WC cells demonstrate apparent blebby morphology in the rounded state. **E**. Polarized WC cell appears with a smooth surface (left and right images) while the rounded cell is actively blebbing (center) until a new protrusion appears (movie 11). The correlated kymograph (10 min interval) presented below each image taken at the position shown on E. **F**. Fluorescence images of WC cell with Lifeact-GFP expression demonstrate the appearance of blebby surface morphology only on the side of protrusion which partially retracted (movie 12). **G**. The blebbing that normally occurs during retraction (red arrows) may subside if another protrusion is extending at the same time. Scale bars A= 30 μm, B,C E= 20 μm, D= 5 μm, F=10 μm.

Next, we investigated the presence of CSE on the surface of the highly aggressive and motile MDA-MB-231 human breast cancer cells (Figure 5A). High resolution fluorescence imaging shows that similar to the CHO and WC cells investigated before, rounded MDA-MB cells in 3D collagen possess a highly convoluted surface morphology with many blebs (Figure 5B). After polarization, these cells develop strong attachments to collagen fibers and are capable of substantial matrix modification (Figure 5D). Polarized cells in 3D collagen have elongated shapes with well-defined long actin cables around the entire cell body beneath the plasma membrane and, in contrast to rounded cells, exhibit little or no apparent surface blebbing (Figure 5C,E). We assume that many blebs or bulges, often visible at the end of the extending protrusions in mesenchymal cells, serve as a membrane reservoir for protrusion growth. An example of how a large bleb on an MDA-MB-231 cell decreases in size simultaneously with the protrusion extension is presented in Figure 5F and <u>movie 13</u>.

**Figure 5.**
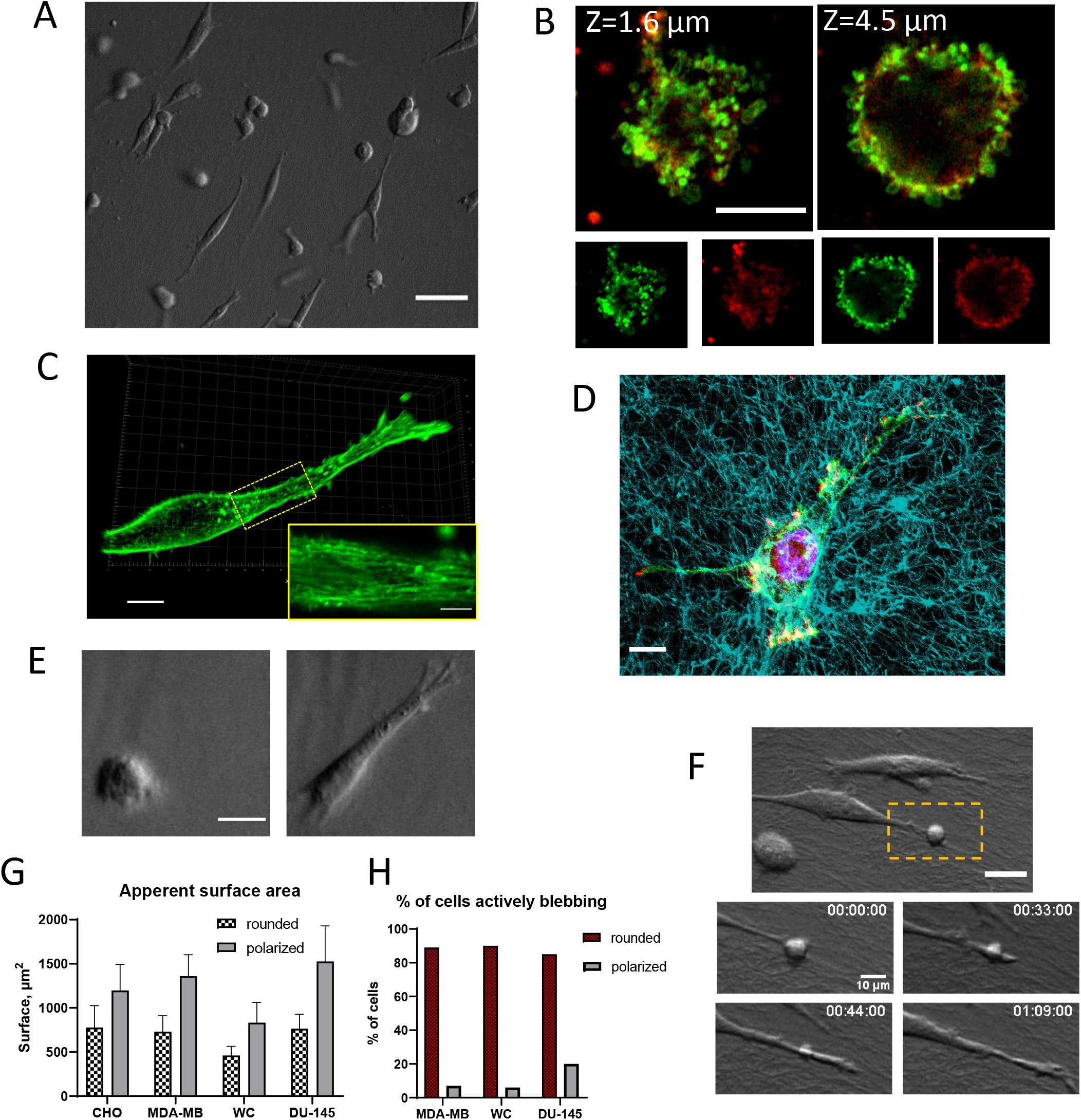
Correlation between protrusion dynamics and apparent blebbing in MDA-MB-231 cells. **A-E.** MDA-MB cells in 3D collagen. **A**. DIC image demonstrates collagen modification and alignment by MDA-MB cells in 3D collagen. **B**. Airyscan high-resolution fluorescence images of rounded MDA-MB cell fixed and stained with Phalloidin (green) and membrane dye (red). Merged (top) and single channels (bottom) images depict the small structures on the cell surface at two different positions along the cell body. The left image, (Z=1.6 um) shows the invasive small protrusions facing out of the cell body into the collagen matrix. Note the perfect colocalization of F-actin and membrane signals which provide evidence that the F-actin signal originated from surface projections. **C.** 3D reconstruction of polarized MDA-MB cell in collagen. Insert on K (magnified part depicted by yellow rectangle, scale bar 5um) presents a magnified image of a focal plane near the ventral part of the cell, depicted by the yellow box. The image shows an absence of small projections on the cell surface and the presence of long actin fibers around the cell body near the interface with collagen fibers. **D**. Collagen modification and alignment induced by MDA-MB cells. Merge confocal image of fluorescently labeled collagen (cyan) and MIP image of F-actin (green), vinculin (red), and nucleus (magenta). **E**. DIC image of representative MDA-MB cell in collagen shows the transformation from a rounded cell with apparent blebbing into a polarized cell with a smooth surface. **F**. MDA-MB can reuse the material from the bleb for protrusion growth. Several time steps of the enlarged area (yellow rectangle) demonstrate how bleb is vanishing simultaneously with the protrusion elongation. (movie 13). **G**. The chart presents the comparison between the apparent surface area of rounded and polarized cells for different cell lines embedded in collagen. The average values were derived for CHO, MDA-MB, WC, and DU-145 cells using

We next investigated DU-145 cells (SupFigure 3A), which in 3D collagen remained mostly stationary with rare, random translocations for the entire duration of the recordings (>24h). After embedding into the collagen matrix, a rounded DU-145 cell with a large amount of stored CSE (SupFigure 3D,E), typically initiates multiple short protrusions from blebs around the periphery (SupFigure 3B) similar to those we observed on SEM micrographs (Figure 2G). After polarization, the blebbing activity disappears around the cell body and mostly remains apparent only on the edges of the protrusions (SupFigure 3C). Polarized DU-145 cells in 3D collagen have mostly irregular shapes with many short protrusions and smooth surfaces as evident from high-resolution fluorescence microscopy (SupFigure 3G). Some elongated cells revealed well-defined long actin cables around the entire cell body beneath the plasma membrane similar to MDA-MB cells (SupFigure 3H,I).

Using DIC images of cells in collagen, we estimated and compared the apparent surface area for the WC, MDA-MB, and DU-145 cells when they were in the rounded state with the area of polarized cells. Our estimates show that the apparent area of polarized WC was up to 3 times larger than the apparent surface area of rounded WC cells (the average ratio was 1.8, N=28). The apparent surface of the polarized MDA-MB cell was 1.8 times larger than the apparent surface of a rounded cell, and it was 2 times larger for DU-145 cells (Figure 5G). The analysis of our video recordings again confirmed a positive correlation between apparent surface blebbing and rounded cell shape for mesenchymal cells in our experiments (Figure 5H).

Many mesenchymal cells embedded in collagen, similar to CHO cells, exhibit oscillatory behavior. Oscillations often arose after the retraction of a long protrusion, which is analogous to the initiation of oscillations after cell rounding from the spread state. This fact provides additional evidence that cells in 3D collagen maintain a large amount of CSE that can be used for rapid shape transformations.

### 6. Morphology of mesenchymal cells without intact microtubules in 3D collagen

Prior studies have shown that the protrusion development and 3D migration of mesenchymal cells, in contrast to 2D migration, is highly dependent on microtubules, although the exact role of MT is not completely understood [40–42]. In our previous investigations of cell oscillations [19, 43], we demonstrated that the amplitude and speed of cell morphological changes are substantially amplified by MT depolymerization, which implies a role of MT in the regulation of CSE. Therefore, we next evaluated the effect of MT depolymerization on changes in shape and surface dynamics in mesenchymal cells in 3D collagen.

All three cell lines with the mesenchymal type of motility used in our experiments after polarization in collagen demonstrated the presence of MT filaments throughout the entire length of the protrusions, even extremely long protrusions such as those in WC cells (SupFigure 4A-D).

When a MT depolymerizing drug was added at the time of embedding cells, all investigated cell types that normally exhibit mesenchymal motility (WC, MDA-MB, and DU145) remained stationary with rounded morphology and were unable to develop long and stable protrusions (SupFigure 5A). However, short protrusions (~5-10 μm) were still able to randomly emerge from the cell periphery. These short protrusions were highly dynamic and were unable to form a stable shape with a smooth surface. We found that similar to the mesenchymal cells, microtubule depolymerization in CHO cells prevented the formation of stable lobopodial protrusions (SupFigure 5B,C). However, without MT, CHO cells were still able to spontaneously translocate in an amoeboid fashion.

### 7. Cell surface excess during amoeboid motility of CHO cells

The amoeboid type of motility, despite the general name, has at least two very distinct modalities [44]. The so-called “blebby-type” amoeboid motility occurs when a cell develops protrusive bulges that push through interstices in the extracellular matrix. The other type of amoeboid motility, which is prominent in leukocytes, is characterized by protrusive short sheets of lamellipodium for navigation in the extracellular environment [26, 44–46].

To investigate the magnitude of the changes in surface area during blebby-type amoeboid motility, we studied the morphology and behavior of the sub-population of CHO cells that employs this type of migration (Figure 6A and <u>movie 2 and 14</u>). We noted above that blebby motility shares many similarities with the compression-dilation mode seen in oscillatory cells (movie *2-4),* where a part of a convoluted cell surface stretched and extended during morphological oscillations in collagen (Figure 6B and SupFigure 1C,D). We postulate that the same mechanism allows a cell to extend, squeeze and eventually translocate through the collagen fibers (Figure 6C, D).

**Figure 6.**
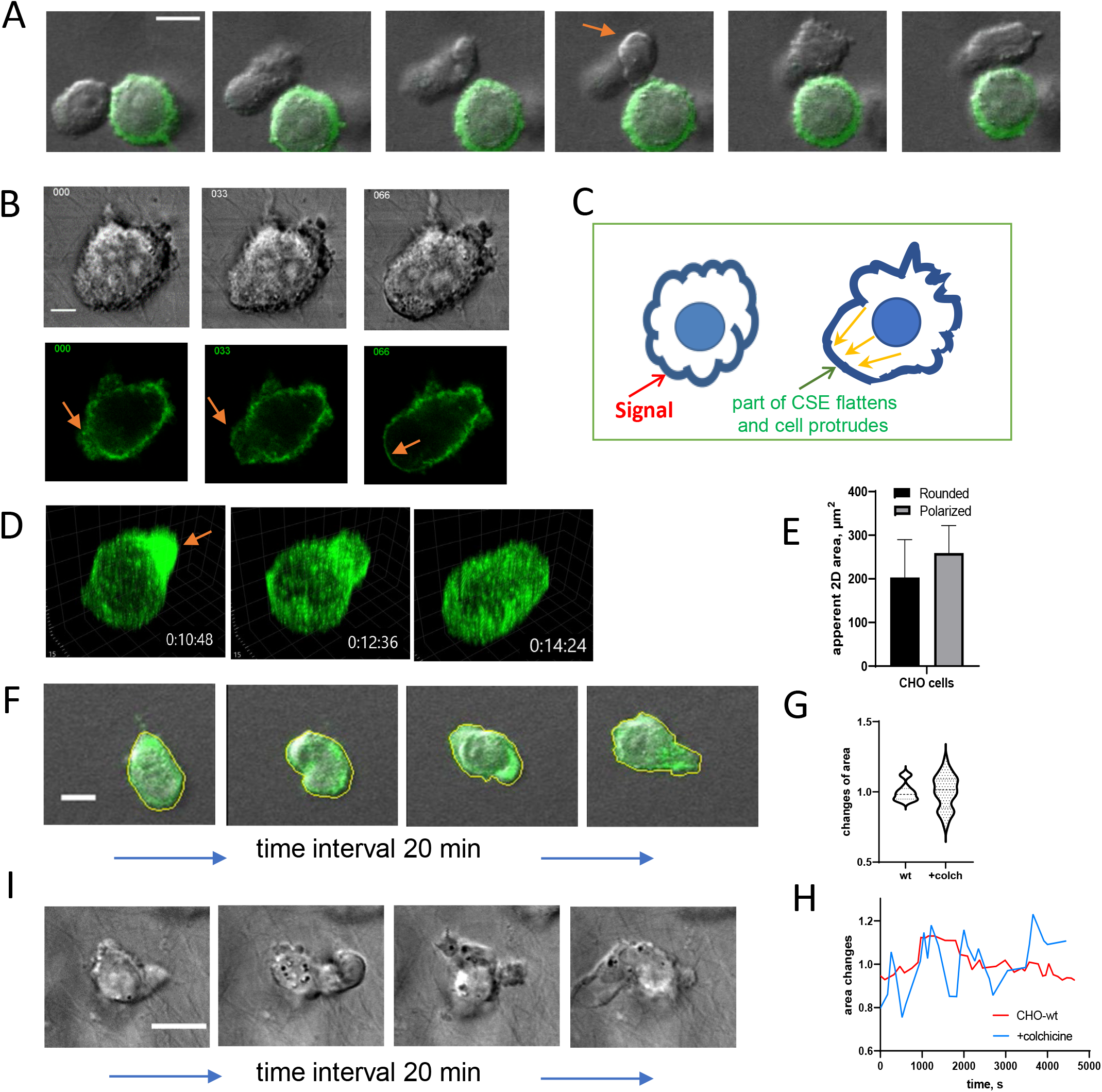
Morphological changes and CSE in CHO cells during blebby-type amoeboid motility. A. Video recording of two CHO cells in 3D collagen. The top cell is translocating using the amoeboid type of motility (movie 2). While pushing through the collagen fibers, the surface of the protruding bulge appears fully stretched and smooth (arrow) for a short period of time. The second cell with an F-actin fluorescence signal demonstrates blebbing and oscillations. B. DIC and F-actin Fluorescence images show stretching/unfolding of the cell cortex that leads to the development of rounded protrusions (arrows). C. The proposed mechanism of cell protrusion. The signal received by the extracellular receptor triggers small projection and folds flattening and CSE release which locally enlarges the cell surface creating additional volume. This volume is immediately filled with cytosol promoting new protrusion. To preserve the total cell volume the other parts of the cell periphery shrink adding more CSE on their surface. D. 3D reconstruction of F-actin fluorescence signal from Z-stack of confocal images (movie 15) of CHO cell during amoeboid motility in collagen demonstrates protrusion in the position with high actin density (arrow). E. Comparison of average 2D area values between rounded cells (N=120) and polarized cells with amoeboid movement (N=10 cells, 20 time frames for each, 3 experiments). F. Morphological changes with the example of area segmentation of CHO cell during amoeboid motility (movie 14). G. Changes in 2D area of CHO cells before and after MT were depolymerized. Values were normalized on the average value for each group (N=10 cells). H. Time plot of 2D area changes for two individual cells with and without MT. I. Examples of shape dynamics of the CHO cell with depolymerized microtubules (movie 16). Scale bars: A,F,I = 10 μm, B = 5 μm.

During amoeboid movement, the CHO cells become more elongated, with an average circularity of 0.81. The complicated cell shape during their motility made it difficult to calculate the apparent surface area from the DIC signal. Therefore, we used as a proxy for these changes the apparent 2D area (XY) measured by segmentation of the periphery near the equatorial section of cells fully visible in the focal plane (Figure 6F). We found that the difference in average 2D area of an equatorial section between rounded and polarized cells is around 20% (S_2D_pol_=245± 63 μm^2^ N=24 vs S_2D_rounded_ = 203 ±86 N=126 for rounded cells) while changes of an individual cell during amoeboid motility are below 14% (with the average 5.8% ± 4.1%, N=6). The visual evaluation showed that during amoeboid motility, cells always have high surface dynamics with apparent blebbing. Similar to an oscillating cell, a large rounded protrusion that eventually can lead to cell translocation appears on the cell periphery mostly in or near the position with a large amount of CSE visually evident by the increased actin density (Figure 6D, <u>movie 15</u>). This large spherical protrusion during a short period of extension (Figure 6A-D) appeared smooth and devoid of blebbing.

While the structure and function of MT in mesenchymal cells were studied intensively, especially on 2D substrate [47–50], much less is known about the role of MTs during amoeboid migration [41, 42]. In our experiments we found that, although MT depolymerization in CHO cells did not completely abrogate the random amoeboid translocation of these cells, the morphological changes became erratic, demonstrating faster dynamics with greater amplitude (Figure 6G,H,I <u>movie 16</u>) This result is consistent with the effect on the oscillating cells on a solid substrate after MT depolymerization [43].

### 8. Cell surface excess during amoeboid motility of leukocyte type cells

To study the dynamics of cell shape and associated changes of surface area in cells with a type of amoeboid migration observed in leukocytes [44, 46], we used U937 cells, a human hematopoietic cell line (monocytic). Similar to CHO cells with amoeboid motility, the surface of U937 cells embedded in collagen was highly dynamic. The presence of blebs and small fin-like surface projections was also confirmed by fluorescence imaging of U937 cells in the collagen matrix (SupFig5 A-D). The rounded U937 cells have an average apparent radius of R=7±0.7 μm. After several hours in collagen, a sub-set of cells becomes spontaneously polarized and motile (Figure 7A and <u>movie 17</u>). The motile cells have a slightly elongated or irregular shape (average circularity index of 0.78±0.1) with an average speed of 0.98±0.70 μm/min, N=20. The measurement of 2D area during motility shows that the difference between rounded and polarized cells during migration on average is below 25% (Figure 7B).

**Figure 7.**
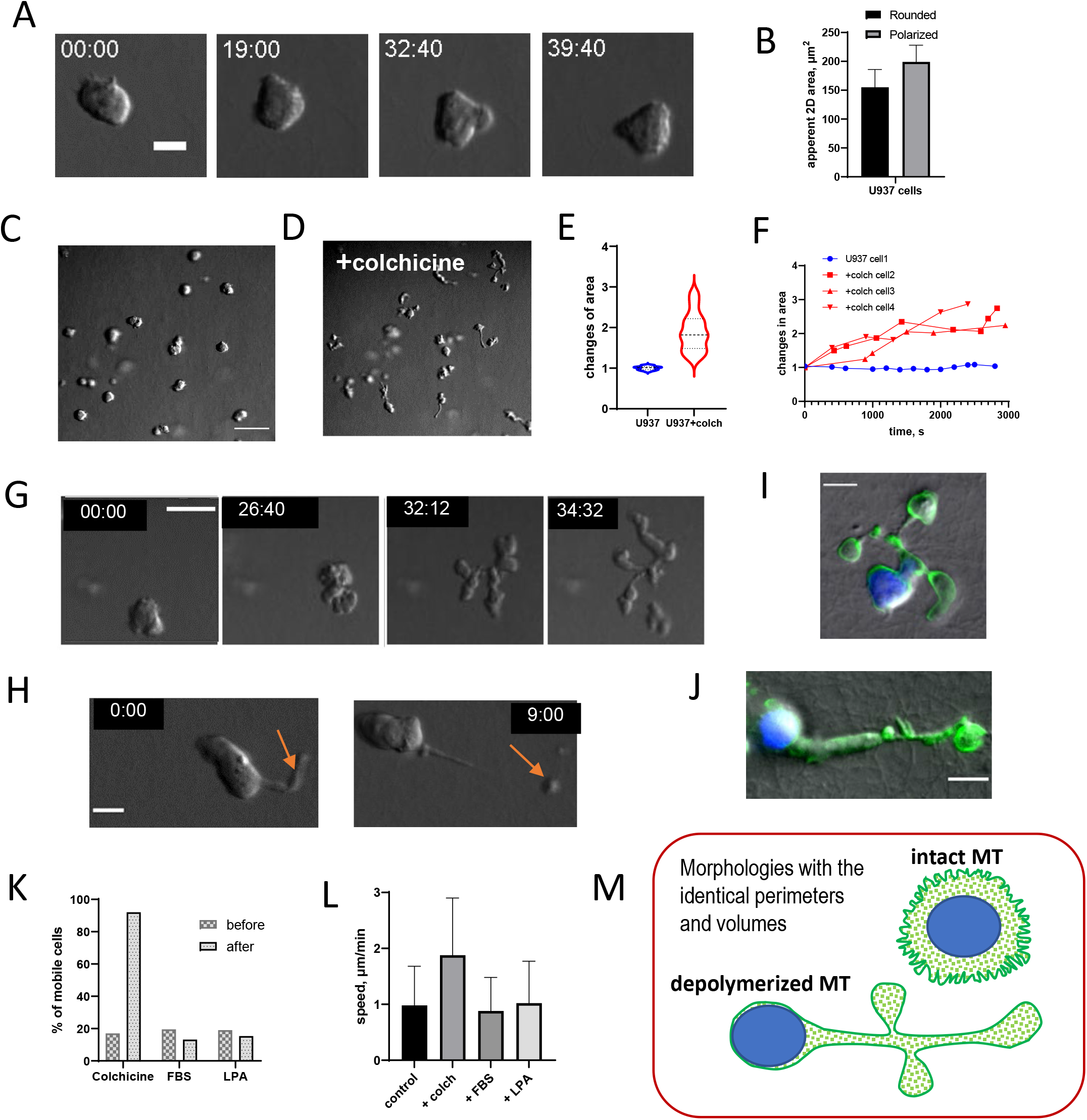
U937 cells motility in 3D collagen and effect of MT depolymerization. A. Time steps of U937 cell shape transformation during motility in 3D collagen (movie 18). B. Average 2D area of rounded and polarized U937 cells in the collagen. C,D. DIC image of U-937 cells embedded in 3D collagen for 24 h with intact MT system (C) and the same cells in 30 min after addition of 1 μM of colchicine (D) (movies 17 and 18). E. Average normalized 2D area of U937 cells during migration with intact MT and after one hour with depolymerized MT. F. Trajectory of normalized 2D area changes after MT depolymerization for three different U937 cells in comparison to area changes for the cell with intact MT. G. Shape transformation of U937 cell after a colchicine treatment (at t=0 s). H. DIC record shows U937 cell with depolymerized MT which has an elongated shape with a long tail (arrow). The tail often gets stuck between collagen fibers, hindering cell movement until it breaks away from the cell (movie 19). I,J, Two merge DIC and MIP of fluorescence confocal images (Phalloidin) of U937 morphology in 3D collagen after MT depolymerization demonstrate smooth surface without blebs. K Percentage of mobile U937 cells and (L) average speed of cell random motility in 3D collagen under different treatments. M. Cartoon illustrates our hypothesis of how rapid release of stored CSE after MT depolymerization transforms cell morphology while maintaining stable cell volume.

MT depolymerization in U937 cells embedded in collagen (Figure 7C,D.) resulted in substantial changes in cell morphology, surface dynamics, and speed within 30 min post-application of the depolymerizing agent (movie 18). Around 90% of cells transformed from rounded or slightly elongated shapes into a morphology that combined bulges of 3 to 5 μm in diameter with long tubules up to ~30 μm in length (Figure 7G). Fluorescence imaging of F-actin showed that a layer of actin cortex underlaid the cell surface while the small surface projections, including blebs, almost completely vanished after MT depolymerization (Figure 7I,J). The abrupt disappearance or flattening of numerous small surface structures inevitably resulted in an abundance of surface area that was stored in them. Our measurements of the 2D cell surface area of some individual cells show increases of up to 300% 30 min after the addition of colchicine (Figure 7E,F). This sudden release of CSE appears to force cells to adopt the morphology of long tubules allow for the accommodation of this CSE without using the surface projections storage (cartoon Figure 7M).

Concomitant with the changes in morphology after MT depolymerization, the percentage of U937 cells randomly moving inside the collagen matrix increased markedly up to 92% and the cells almost doubled in speed (average speed 1.9 ± μm/min). The changes in cell morphology and dynamics persisted for the time of experimental recording (up to 10h post colchicine treatment).

The fast moving cells often adopted a long tubular shape (Figure 7H, <u>movie 19</u>), with which these cells appeared to “flow” through the matrix. These cells also frequently acquired long tails that were often enmeshed in the collagen fibers constraining cell movement until the tails broke off. We believe that such tails appeared in the absence of intact microtubules as a collector for CSE that can no be longer distributed around the cell body.

It was established previously that the amoeboid migration is dependent on the Rho/ROCK signaling pathway [32, 51], which regulates the contractility of the acto-myosin cortex. The Rho guanine nucleotide exchange factor, GEF-H1, is the known mediator that connects RhoA activation, actin polymerization, and myosin contractility to microtubule dynamics. GEF-H1 is inactive when it is sequestered by MT and upon release from MT becomes active [52, 53]. Therefore, the MT depolymerization sharply increases the level of active GEF-H1 followed by the increase of RhoA and Myosin II contractility. To test whether the strong effect of MT depolymerization on the shape and motility of U937 can be solely attributed to increased myosin II contractility via the GEF-H1-RhoA-ROCK pathway, we stimulated the activity of RhoA and myosin II by 30% FBS or 1μm LPA treatments after cell starvation. However, neither treatment led to shape changes, membrane tubules, an increase in the number of migrating cells, or migration speed (Figure 7K,L, SupFigure 5G,J).

## Discussion

Cells require a reserve of surface area in order to successfully execute fast shape transformations without threatening plasma membrane integrity. Despite its importance, the regulation of cell surface area has not attracted as much attention as the regulation of cell volume or membrane tension since it has been assumed that the surface area is a passive partner in changes of cell shape, simply following the morphology dictated by membrane tension and the cytoskeleton. However, cells generally have much more surface than is required to simply cover the cell volume, implying that specific mechanisms of surface area regulation should exist in addition to endo-exocytosis. We suggest that the simplest and most efficient mechanism for surface area regulation is a controlled development of small surface projections: blebs, ruffles, filopodia, microvilli, etc. These projections can store massive amounts of cell surface and are highly dynamic, which means they can be rapidly dismounted with the release of local CSE and immediately rebuilt in other locations on the cell periphery.

Our investigation of cell dynamics during 3D migration in collagen revealed that cycles of high cell surface activity (blebbing and oscillations) were followed by quiescence and visible surface smoothing. Similar cycling behavior between protruding and migrating cells (“running”), and more static cells that remain in the same position but with a loss of cell polarity (“tumbling”), was found during the migration of primordial germ cells [54–57], mammary epithelial cells [58], invasive metastatic cells [59, 60] and during bacterial chemotaxis [61]. It was suggested that this running and tumbling behavior represents an intrinsic property of migrating cells[62]; however, the mechanism underlying this behavior was not established.

Our analysis shows that intervals of surface smoothing on the cell body correspond with the extension of large protrusions, while active blebbing corresponds to cell rounding after the withdrawal of large protrusions. Fluorescence analysis of mesenchymal cells in collagen matrices revealed that while rounded cells have a large number of blebs and other surface projections, polarized cells with long protrusions have substantially smoother surfaces, which is consistent with our hypothesis that cells release CSE stored around the cell body to cover the newly developing protrusion. During the protrusion retraction, the resulting CSE requires rapid packing, which is achievable by forming new blebs as they present the fastest attainable form of storage.

For the past two decades, blebs have often been considered to be an artifact arising from a compromised connection between cortex and plasma membrane which is pushed out by high pressure in the cytosol [63, 64]. It should also be noted that the majority of information about blebs has come from studies of cells spread on a solid 2D substrate or during artificially induced blebbing [65–73]. Moreover, for purposes such as the estimation of cortex thickness or modeling the biomechanical properties [73–77] it was often assumed that the membrane and actin cortex are two perfectly smooth concentric layers with uniform density. However, bearing in mind that the plasma membrane is essentially not stretchable, under this assumption the process of blebbing would not be possible because in order to develop one typical bleb with a radius of r=1 μm, an additional plasma membrane area of ~12 μm^2^ is required. The fact that the average time of bleb expansion is less than 30 sec leads us to the conclusion that the additional surface must come from some readily available source in the close vicinity of the growing bleb. Cell rounding or protrusion withdrawal each generate large amounts of CSE that creates a favorable condition for numerous blebs production. For the cells with stable morphology, the most obvious source for new bleb is a shrinking bleb or filopodium nearby which frees surface area for reuse. Several studies already confirmed that the probability of a new bleb appearance is higher around the position where the other bleb is shrinking or after the disappearance of filopodia [36, 78, 79].

In this paper we emphasized that surface blebs provide an efficient mechanism for prompt storing of surface excess. Because a typical lifetime of a single bleb is 1~ 3 minutes this mechanism also allows the rapid redistribution of surface area and therefore should be considered as an important part of normal cell physiology with the specific regulatory pathway which needs to be established.

Our data indicate that MT play an essential role in CSE regulation and stability. We hypothesize that the intact MT system coordinates cell surface area with cell volume by regulating the assembly and release of CSE. We also assume that MT can stabilize cell shape by restricting CSE dynamics. By contrast, MT depolymerization prevents long-range coordination of CSE regulation, which leads to uncontrolled excess of cell surface which, in turn, accelerates shape changes. Consistent with this hypothesis, it was reported that tethers extracted from cell membranes after MT depolymerization became significantly longer than in control cells and even longer than in cells with depolymerized actin filaments [80–82]. Similarly, micropipette aspiration studies of living cells revealed substantial cell softening after MT depolymerization [83, 84].

Our hypothesis can also explain the puzzling observation from previous studies that MT depolymerization blocks mesenchymal cell movement in 3D while enhancing amoeboid motility [41, 85]. We show here that despite many differences between mesenchymal and amoeboid migration, both types rely on cell surface stored in small surface projections. Because the morphological changes during mesenchymal migration are slow and the shape changes during the amoeboid type of translocation are rapid, these two modes require the deployment of CSE within different time scales. For mesenchymal cells, stable shape and slow surface dynamics are important factors for building protrusions, developing adhesions, and transforming the ECM by enzymatic degradation. We hypothesis that dysregulation of CSE dynamics after MT depolymerization prevents the delivery of the surface from the cell body CSE which is necessary to cover the growing protrusions. In the absence of a coordinated supply, cells are only able to extend short protrusions using the cell surface locally available on the periphery. Furthermore, without the shape stabilizing function of microtubules, these protrusions are likely to collapse shortly after development. We believe that during 2D mesenchymal migration, MT are dispensable because strong adhesions to the substrate stabilize the shape and diminish the role of MT.

In contrast to mesenchymal motility, during amoeboid migration, the rapid shape transformation necessary for squeezing between ECM filaments requires fast CSE dynamics. For fast-moving cells undergoing ameboid motility with short protrusions, local reserves of CSE are enough for providing the needed cell surface. In this case, the MT network stabilization function is important for restricting CSE release everywhere on the cell periphery except for localized release in the direction of movement. This locally released CSE is inflated by cytosol and followed by actin polymerization inside the protrusion and stabilization by newly grown MT. When MT are depolymerized, CSE is released without spatial regulation, forcing cells to adopt a new morphology, creating conditions for rapid actin polymerization in different directions, leading in some cases to cell fragmentation. Therefore, MT control of CSE regulates the cell protrusion, retraction, and stabilization required for dynamic cell movements and migration.

## Methods

CHO-K1, MDA-MB-231, DU-145, and U937 cells were obtained from the Tissue Culture Facility of *Lineberger Comprehensive Cancer Center*, UNC at Chapel Hill. WC cells were acquired from the ATCC (Manassas, VA). CHO cells stably expressing Lifeact-GFP (the small 17-aa peptide, Lifeact, fused to GFP) and expressing myosin regulatory light chain fused to RFP (MLC-RFP) were a gift from the James Bear laboratory (UNC at Chapel Hill). MDA-MB and DU-145 cells were cultured in DMED (Gibco) media with 10% FBS (Gibco). CHO cells were grown in DMEM/F12 (Gibco) media containing 10% FBS and 4 mM L-glutamine. WC and U937 cells were grown in media RTMI (Gibco) containing 10% FBS. All media contained 100 U/ml penicillin/streptomycin. Colchicine (Sigma-Aldrich), LPA (Sigma-Aldrich), FBS (Gibco) were used for cell treatment during the experiments.

### Electron microscopy

The samples for SEM and TEM were prepared according to the protocol published previously [20, 33]

### Collagen preparation

Bovine collagen I solution from Advance Bio Matrix: PureCol (3mg/ml, #5005) or PureCol^®^ EZ Gel (5mg/ml, #5074) were used for experiments. The 0.5 M NOH was added to adjust the pH level in the case of PureCol solution. For the experiments, we used glass bottom dishes, 14mm glass, N=0 from Mattek or CellVic companies. The thin glass (N=0) allowed visualizing a thicker slice of the collagen matrix. All gel preparations were carried out on a table cooler located inside a tissue culture hood. To ensure that experimental cells were located inside the collagen matrix and were far from the hard substrate, we embedded cells in a collagen “sandwich”.

For the first layer of the sandwich we used 40 μl of cold collagen solution adjusted appropriately for the cell line medium to the concentration used in the experiment. This solution was distributed evenly on the bottom glass and allowed to polymerize for 10 min in the incubator. Then 100 μl of collagen solution with the cells (~10^6^ /ml) were placed on top of the first layer and polymerized for 1h in the incubator. After 1 h, 500 μl of media was added to the top of the collagen sandwich and the dish was either kept in the incubator or used for recording.

For visualization of collagen structure 10% of collagen volume was substituted by collagen-FITC (Sigma-Aldrich) and used for creating the collagen matrix.

### Collagen fixation and cell labeling

The collagen matrix with cells was fixed using 2ml of warm (37C) paraformaldehyde solution [4% (w/v) in PBS, pH 7.4] for 20 minutes, washed twice with PBS, and incubated with blocking/permeabilization solution containing 5% goat serum, 1% BSA, and 0.1% saponin in PBS for 1 h at room temperature. Samples were incubated overnight at 4C with primary antibodies following 3x washing and incubation for 1h at room temperature with appropriate secondary antibodies.

All primary antibodies and their matching species-specific Alexa fluor conjugated secondary antibodies were diluted in PBS supplemented with 1% BSA and 0.1% saponin. Excess antibodies were removed by three washes with PBS containing 0.05% saponin, and the nuclei were counterstained with Hoechst 33342 (1:2000) (Invitrogen) for 8 mins at room temperature. After washes with PBS containing 0.075% saponin, collagen with cells were washed with PBS, covered in 1ml PBS, and directly visualized under a confocal microscope as described below. Collagen fixation, labeling, and washing were all performed extremely gently to avoid perturbing the collagen structure or detaching the matrix from the glass.

Primary antibodies used in the experiments: Anti-vinculin (Sigma-Aldrich, SAB4200729), Antitubulin (Santa Cruz Biotechnology). To visualize F-actin in fixed cells we used Alexa Fluor™ 488 or 568 Phalloidin (Invitrogen™); Species-specific Alexa-fluor conjugated secondary antibodies were purchased from Thermo Fisher Scientific. For PM visualization in live cells, we transiently expressed the PMT-mRFP which is a lipid-linked protein that resides almost exclusively on the inner leaflet of the PM and described elsewhere [86]; The PM in fixed cells was stained by CellTracker™ Deep Red Dye (Invitrogen)

### Microscope imaging

The long-term time-lapses in DIC mode were recorded on the Olympus VivaView incubator microscope which provides the ability to record up to 8 samples simultaneously with full control over position and acquisition parameters. The images with fluorescence signal of cells in the collagen were recorded with a 40x silicon immersion objective using an Olympus FluoView1200 laser scanning confocal microscope with an environmental chamber. The super-resolution images were acquired on Zeiss 880 microscope with Airyscan detector using 40x oil objective.

### Image and statistical analysis

Image analysis was performed using ImageJ software. The cell speed was estimated using the Manual Tracking plugin from ImageJ.

Statistical tests were performed using Prism 7 software. The statistical tests applied to each quantification are indicated in each figure legend. ‘+/- ‘ indicate standard deviations.

### Estimation of surface area

#### For rounded cells

To estimate the apparent surface of rounded cells we measured cell perimeter on the cell equator after manual segmentation. The diameter was calculated as d=perimeter/π and the surface of a rounded cell was calculated a s ***Surface*** = **π**\***d^2^**

#### Cigar-shaped cells

To estimate the surface area of a polarized cell with protrusions we assumed that the shape of the cell body can be approximately presented as a prolate ellipsoid (cigar shape) and be calculated after measurement of its axis: long (c) and short (a).

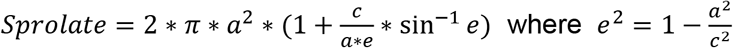

The surface of the protrusion was estimated as a surface of a tube with a diameter measured in the middle of protrusion length. For the measurement, we selected cells with symmetrical, cigar-shaped bodies that were fully visible on the focal plane during at least 10 consecutive frames.

#### Estimation of cell surface for irregularly-shaped DU-145 cell in collagen

Using ImageJ software the cell shape was manually segmented and morphological parameters (area and perimeter) were measured. Cell surface was approximated as twice the visible area (2*Area, apical and ventral surface) plus the length of the perimeter multiplied by 1 μm (1*Perimeter) to approximate the surface area due to cell thickness: S =2*Area + 1*Perimeter

#### 2D area of amoeboid cells

To estimate the area for the amoeboid cells we manually outlined the cell periphery and measured the enclosed area for each time point in which the cell was fully visible in the focal plane.

## Acknowledgments

We dedicate this manuscript to our friend, colleague, and co-author, Dr. Ken Jacobson, who passed away during the preparation of the final drafts of this manuscript. This work was supported by the National Institutes of General Medical Sciences grant R01GM134531 (REC).

## Authors contributions

MK conceived and designed the study; performed experiments, analyzed data, and wrote the manuscript draft with help from Ken Jacobson. DL, JZ performed experiments, analyzed data and edited the manuscript. BW, VW performed experiments and analyzed data. RC provided resources and edited the manuscript.

---

**Supplemental Figure 1.**
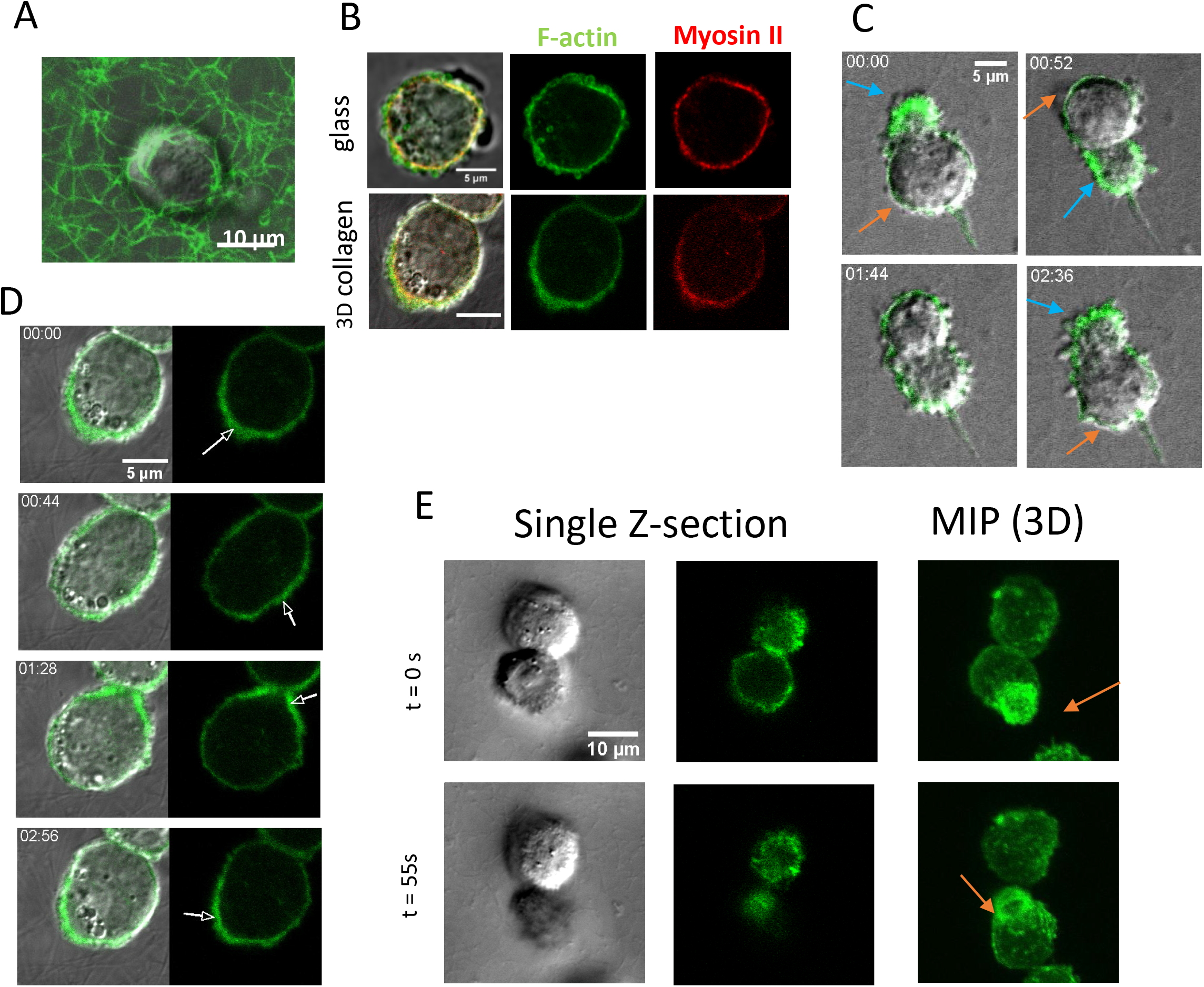
Morphology and oscillations of rounded CHO cells in collagen. A. DIC image of CHO cells embedded in a fluorescently labeled (FITC) matrix shows that collagen fibers are not constrained or modified around the cell which confirms the absence of firm attachment between a cell and fibers. B. DIC and fluorescence images of CHO cells demonstrate the similarity of F-actin (Lifeact-GFP) and myosin II (MRLC-RFP) distributions in the freshly rounded cells on a glass substrate and after 24h in the collagen matrix. C. Example of CHO oscillations inside the 3D collagen (movie 3). Note the correlated changes of cell shape and intensity of Lifeact-GFP signal during one period of oscillation: on the side of expansion, the signal from the actin cortex becomes thinner with apparent small numbers of blebs (orange arrows) while on the opposite shrinking side the signal becomes wider and showing a large number of blebs (blue arrows). D. DIC and fluorescence image of CHO in collagen during one period of oscillation with the apparent traveling wave of cortex density visible as a green patch moving along the cell periphery (arrows), see movie 5. E. DIC and F-actin Fluorescence images (Lifeact-GFP) of CHO cells embedded in the 3D collagen matrix and exhibiting morphological oscillations (movie 4). The top cell oscillates in the XY direction, while the bottom cell oscillates in the Z direction (in and out of the focal plane). Maximum intensity projection images illustrate the smoothing and stretching of the cortex (arrow) during shape extension.

**Supplemental Figure 2.**
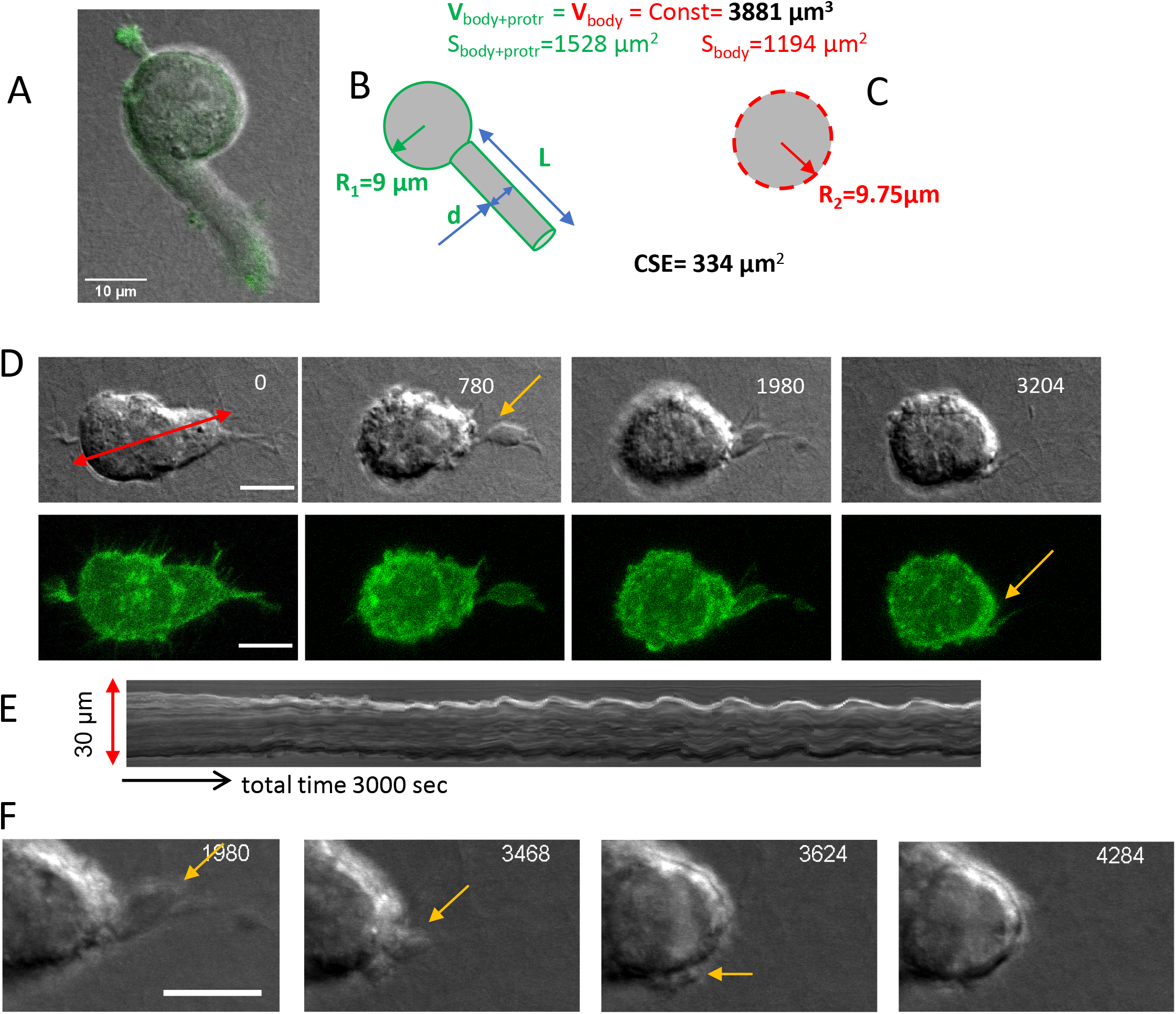
Estimation of CSE in CHO cells with lobopodial protrusion and CSE dynamic after protrusion retraction. A-C. (A). We assumed for simplicity that cell body is a sphere with a smooth surface with the radius R_1_=9 μm and corresponding Volume = 4/3*π*R_1_^3^ = 3052 μm^3^, and surface area S= 4*π*R_1_^2^= 1017 μm^2^. We also assumed that the protrusion has a cylindrical shape with the length L=25 μm and the diameter (2r) measured at the middle of protrusion length equal to 6.5 μm (B). The protrusion surface area can be calculated as Spr= L*π*2r = 510 μm^2^ and the volume as V_pr_=L*π*r^2^=829 μm^3^ The total volume of the cell with the protrusion is V=388l μm^3^ and the total surface area is S=I528 μm2. Assuming that the total volume is conserved, to adopt the additional volume from the protrusion the radius of the cell should expended and became R_2_=9.75 μm (C). However, the increased body size requires only S=1194 μm2 of surface, leaving in excess 334 μm2 of cell surface which is needed to be stored or exocytosed. D. DIC and F-actin (Lifeact-GFP) dynamics during protrusion withdrawal of CHO cell embedded in 3D collagen matrix (time in seconds). E. presents a kymograph at the position shown by red arrow on D. Note the presence of a smooth surface on the cell with extended protrusion, blebbing during the retraction and beginning of oscillations after protrusion retraction (movie 8). Yellow arrow points at the potion of CSE accumulation after protrusion withdrawal. The enlarged part of image D on F demonstrate how the morphological oscillations help to redistribute local CSE around the cell body. Scale bars: A= 10 μm, D=10 μm, F=5 μm.

**Supplemental Figure 3.**
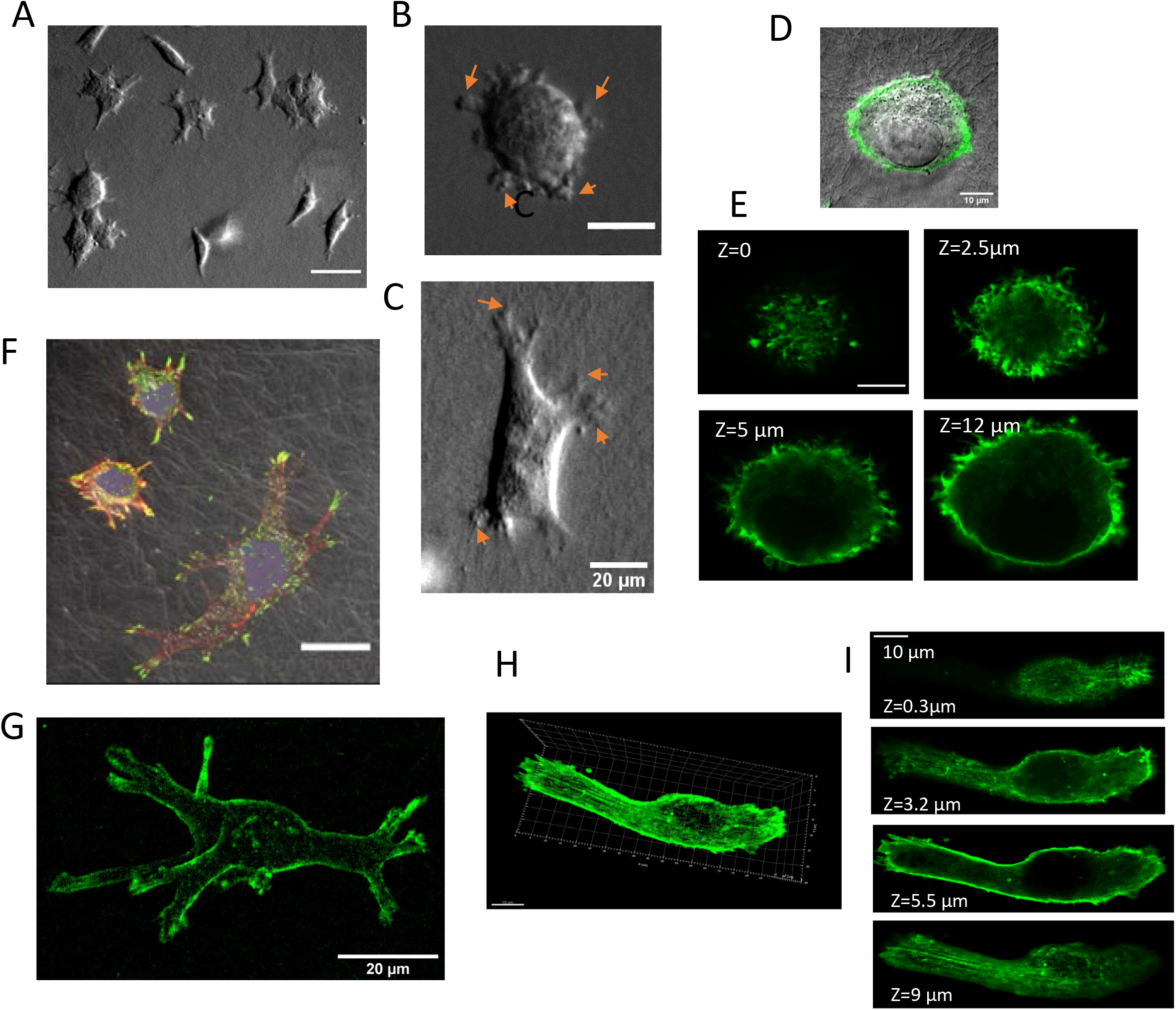
DU-145 cells in collagen I matrix for 24h. A. DIC images depict different morphologies of DU145 cells after 24h in the collagen matrix. B. Initiation of multiple protrusions from the blebs on the periphery of rounded DU-145 cell (arrows), demonstrates similarity to the process depicted on SEM images from Fig.3. C. After cell polarization the blebbing is apparent only on the edges of protrusions (arrows). D,E. The same rounded DU-145 cell in collagen with Phalloidin (green) staining was imaged in two different microscope modes. D. Merged DIC and scanning confocal mode show the F-actin cortex together with the collagen structure. E. High-resolution fluorescence Airyscan image depicts the small structures on the cell surface at the different positions along the cell body. Note how the invasive small protrusions face out of the cell body into the collagen matrix (Z=0). F. Merged DIC and fluorescence 3D maximum intensity projection image (vinculin-green, F-actin-red) of fixed DU-145 cells after 24h in 3D collagen. G. 3D volume reconstruction from Z stack of high-resolution Airyscan images of actin structure (Phalloidin) of polarized cells in the collagen shows that the surface of the cell with multiple protrusions is smooth (reconstructed from Z stack of 75 images. H,I. High-resolution fluorescence Airyscan image data of actin structure (Phalloidin, green) of polarized cell DU-145 in the collagen. 3D reconstruction (H) and separate slices (I) reveal a smooth surface and the presence of long actin fibers around the cell periphery. Scale bars: A=50 μm, B,C,F,G =20 μm, D,I =10 μm

**Supplemental Figure 4.**
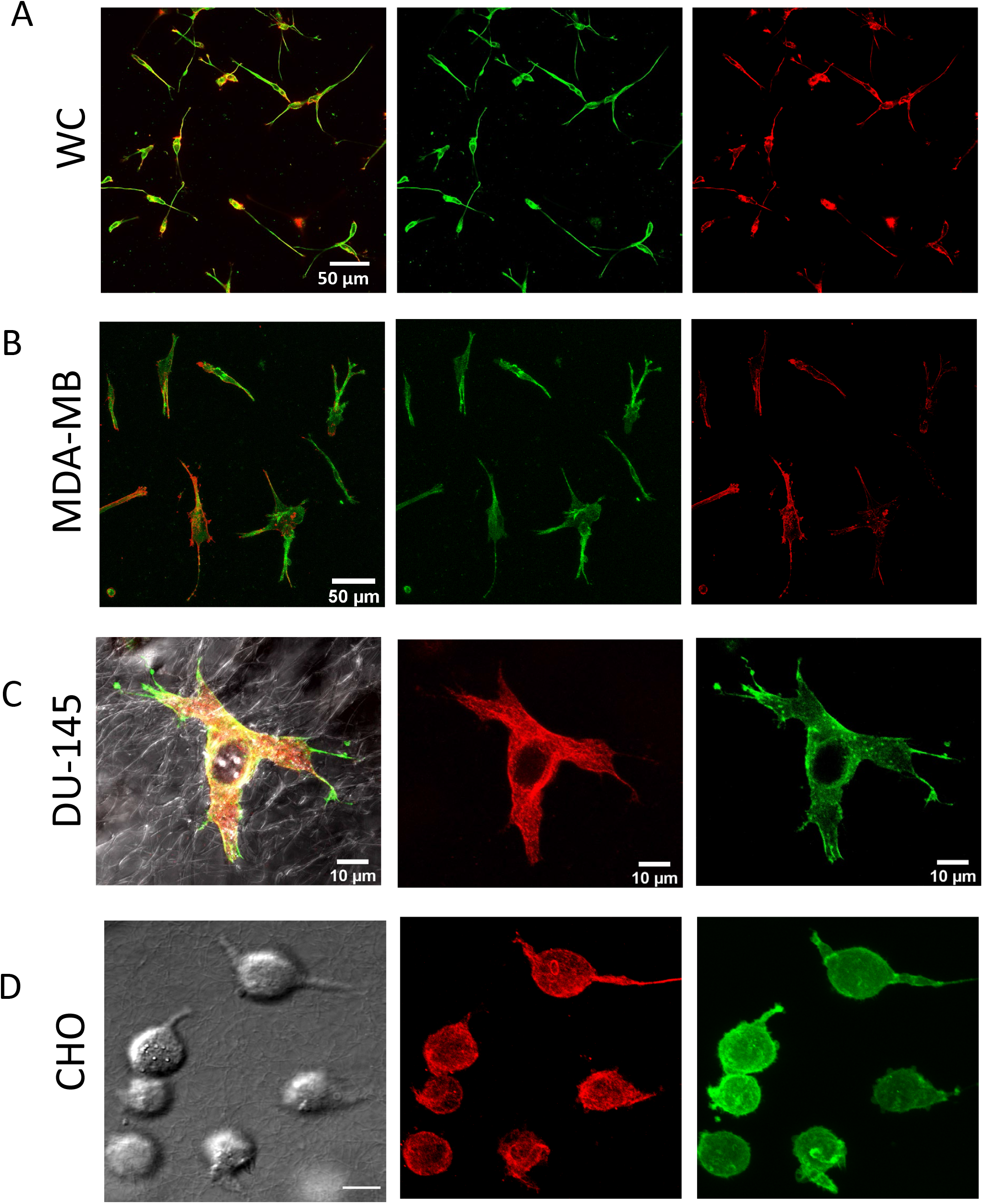
Presence of MT in cell protrusions. A.B. Maximum intensity projection fluorescence images of F-actin (Phalloidin-red) and microtubules (anti-tubulin, green) of WC (A), MDA-MB (B) cells embedded in the 3D collagen for 24 h. C.D. Merged (C) and separate (D) DIC and maximum intensity projection fluorescence images of F-actin (Phalloidin-green) and microtubules (anti-tubulin, red) DU-145 (C), and CHO (D) cells embedded in the 3D collagen for 24 h.

**Supplemental Figure 5.**
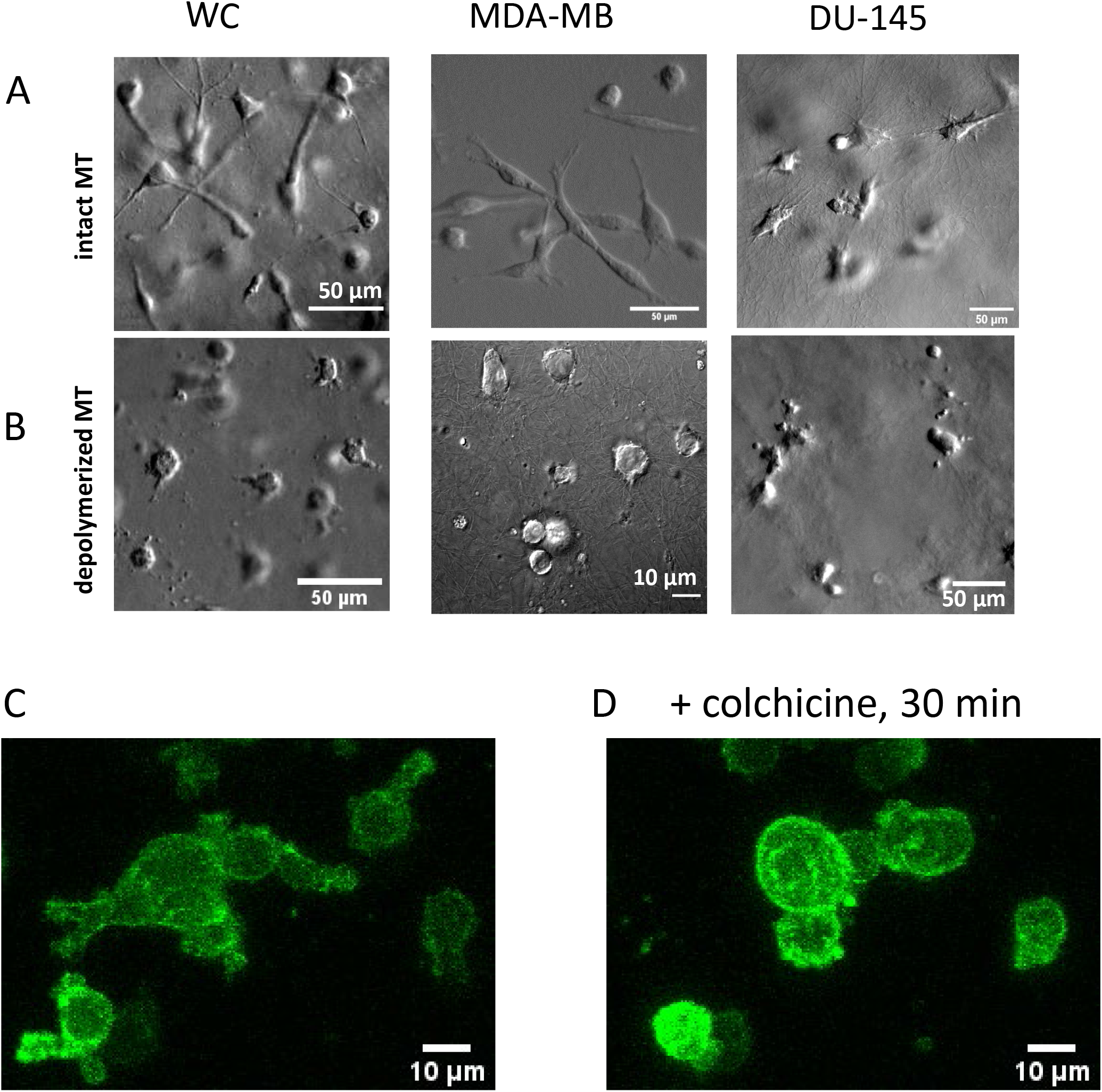
Effect of MT depolymerization on cell protrusions in collagen. A. DIC images of WC, MDA-MB, and DU-145 cells with intact MT system after 24h in the collagen matrix. B. The same cell lines were embedded in the collagen with MT depolymerization agent (1uM of colchicine). C. MIP images of F-actin (Lifeact-GPF) in CHO-cells embedded in 3D collagen for 24 h. D. The same cells 30min after the addition of colchicine. MIP images were built from Z-stack of 61 images (total 60 μm of collagen thickness

**Supplemental Figure. 6.**
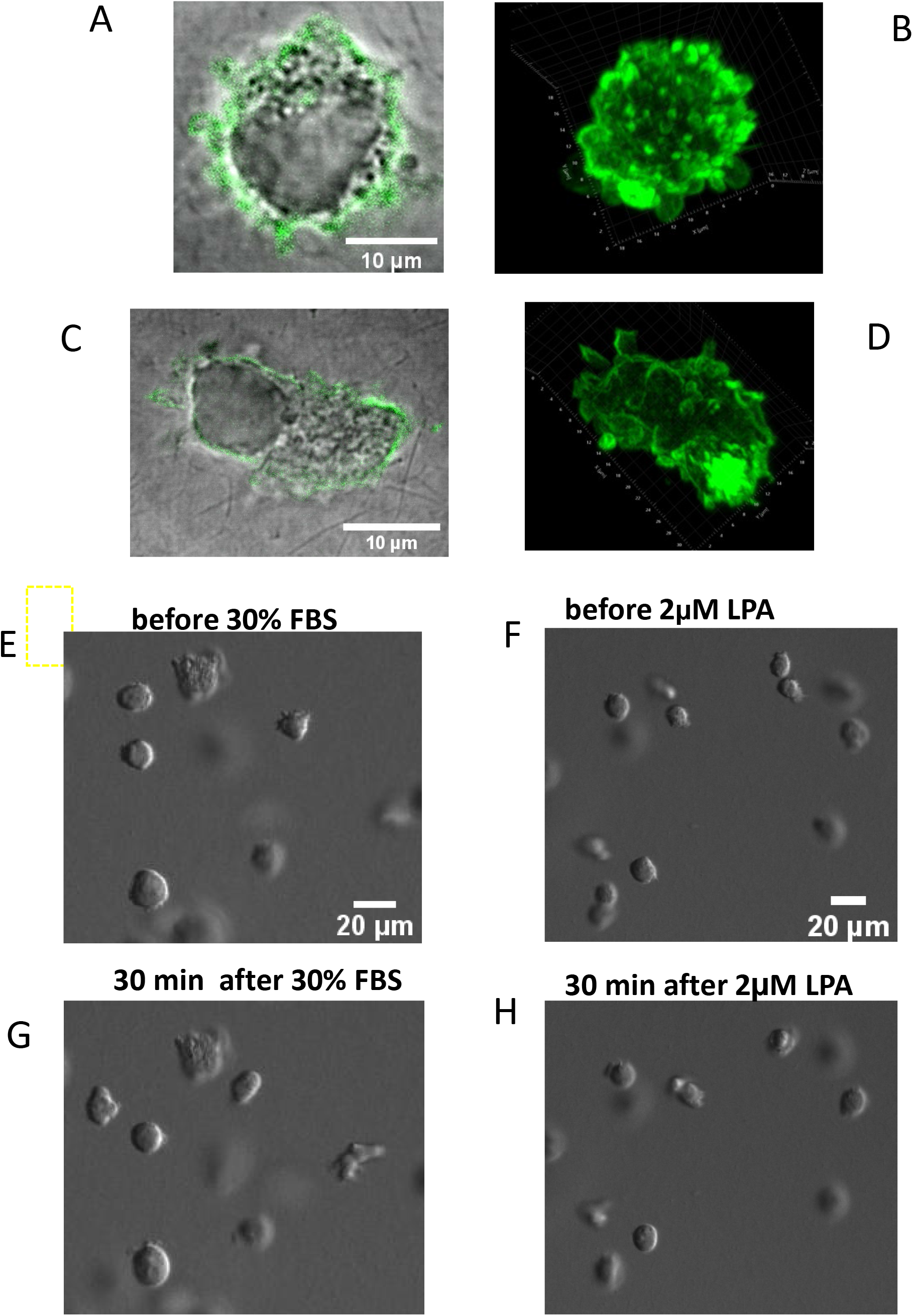
Morphology of U-937 under different treatments. A-D. Morphology of rounded (A,B) and polarized (C,D) U937 cells in 3D collagen stained with Phalloidin (green). A,C. Merged DIC and fluorescence scanning confocal images. B,D. 3D reconstruction of the same cells built from Z-stack of Airyscan high-resolution images showing highly convoluted surface morphology and presence of blebs. E,F DIC images of starved (0.5% FBS) U-937 cells embedded in collagen for 24 h and the same position imaged in 30 min after addition of either 30% FBS (G) or 2 μM LPA (H). Images demonstrate that cell morphology was not affected by either of these treatments.

