## Supplementary material for "Cell surface excess is essential for protrusions and motility in 3D matrix": movie legends

**Legends for Supplemental movies.**

Movie 1. An example of CHO cell migration in 3D collagen using lobopodial type of protrusion.

Movie 2. An example of CHO cell migration in 3D collagen using blebby type amoeboid motility. One cell is moving by developing a bulge and pushing through the collagen while another cell is oscillating without translocation. Green fluorescence signal on stationary cell shows localization of F-actin marker (Lifeact-GFP).

Movie 3. Oscillation of CHO cell after embedding in 3D collagen for more than 24 hours. The F-actin (Lifeact-GFP) fluorescence signal demonstrates stretching of the cell cortex during bulge extension.

Movie 4. CHO cell oscillations in collagen. In the collagen matrix, the periodic cell protrusions during the oscillation can occur in different directions. The top cell oscillates in the XY direction, while the bottom cell oscillates in the Z direction (in and out of the focal plane). The maximum intensity projection image illustrates the unfolding and stretching of the cortex (arrow) during chape transformation in 3D.

Movie 5. The highly periodic process of stretching (dilation) and compression of the membrane cortex layer creates a traveling wave of cortex density around the cell periphery.

Movie 6. The movie demonstrates a smooth surface with fully extended protrusion and the blebbing during and after protrusion withdrawal. Oscillations are initiated at the end of protrusion retraction.

Movie 7. CHO cell in the collagen during phases of tumbles and protrusions. The movie demonstrates a protrusive phenotype of CHO cells in collagen matrices in which these cells undergo morphological oscillations and blebbing prior to the extension of a lobopodial-like protrusion. Simultaneously with the initiation of a new protrusion, the morphological oscillations ceased and the blebbing decreased. After the extension of a large, stable protrusion, the cell exhibited low surface dynamics and only a few blebs.

Movie 8. Correlation between protrusion and blebbing. CHO cell with fluorescence signal from F-actin marker (Lifeact-GFP) appears with a smooth surface while having a big round protrusion. In the initial stages of protrusion withdrawal, the cell surface became covered with dynamic blebs which were highly pronounced after retraction. During the later stages of protrusion retraction, morphological oscillations began. The retraction of protrusion leaves a region on the cell periphery densely populate with blebs that locally store a high amount of CSE. The traveling wave of cortex density during morphological oscillation helps to redistribute this local CSE around the cell.

Movie 9. Walker carcinoma cells demonstrate fast random motility in 3D collagen.

Movie 10. The protrusions and migration of WC cells involve substantial collagen contraction. The movie shows how the WC cell which is located inside the collagen but out of a focal plane is developing two long protrusions (60um and 50 um) that contract the collagen fibers.

Movie 11. The movie demonstrates that WC cells in 3D collagen have substantial blebbing if they acquire a rounded shape during motility. The blebbing on rounded cells ceases simultaneously with protrusion initiation.

Movie 12. Movie of WC cell with the expression of F-marker (Lifeact-GFP) demonstrates the development of blebs only on the retracting part of protrusion.

Movie 13. The movie demonstrates how the MDA-MB cell uses large bleb to extend the protrusion.

Movie 14. Amoeboid motility of CHO cell with Lifeact-GFP fluorescence signal. Video demonstrates that the surface of a motile cell has apparent constant blebbing with local extensions that drive relocations.

Movie 15. 3D reconstruction of F-actin fluorescence signal from Z-stack of confocal images of CHO cell during amoeboid motility in collagen. The movie demonstrates that a large rounded protrusion that leads to cell translocation appears mostly on or near the position with a high amount of CSE visually evident by increased actin density.

Movie-16. CHO cells embedded in collagen with depolymerized microtubule system demonstrate the erratic dynamics of cell shape with faster and greater amplitude changes. The movie shows on the left the Maximum intensity projection of F-actin fluorescence (Lifeact-GFP) imaging and on the right DIC imaging of the same cell.

Movie-17. Example of U937 cells motility in 3D collagen

Movie-18. Effect of microtubule depolymerization on U937 motility. U937 cells were embedded in 3D collagen with intact MT for 6 h before the recording. During the recording, a 1uM of colchicine was added to the media on the top of the collagen matrix.

Movie 19. Example of U937 cell motility in the 3D collagen with depolymerized MT system.
